# Pyramiding four genes governing bacterial blight resistance in three popular rice varieties of East Africa and Madagascar

**DOI:** 10.64898/2026.06.29.735165

**Authors:** Yugander Arra, P.I Eliza Loo, Camille Blasco, Emilie Thomas, BN Devanna, Melissa Stiebner, Mathilde Hutin, Florence Auguy, Boris Szurek, Wolf B. Frommer

## Abstract

Bacterial blight of rice causes substantial crop losses in Asia and Africa. The recent introduction of Asian strains into African countries has led to two independent outbreaks detected in 2019, causing severe damage in Madagascar and Tanzania. The strains are highly virulent on local rice varieties and rapidly spread from Tanzania to neighboring countries. Multiplex genome editing of effector-binding elements in the promoter of SWEET rice promoters has successfully generated elite rice lines with broad-spectrum resistance against bacterial blight. While genome-edited crops can be released in countries with biosafety regulations, their use in countries that have yet to establish regulations, e.g., Tanzania and Madagascar, is hindered. To circumvent this, marker-assisted backcross breeding (MABB) of the African elite varieties Komboka, FARO-44, and NE-RICA-4 was adopted to introgress resistance genes to confer resistance to *Xoo* strains identified in Tanzania (iTz) and Madagascar (iMg). Resistance gene pyramiding, namely *Xa1*, *Xa4*, *xa13*, and *Xa21*, in the three elite rice varieties conferred resistance to iTz and iMg strains. Our study presents a solution to provide rice breeders in East Africa with bacterial blight resistance in three local rice varieties for field trials, line registration, and deployment.

## Introduction

Rice is one of the most important staple food crops. In sub-Saharan Africa, rice has become a major staple food crop in addition to maize, sorghum, and cassava. Over the past decade, the area under rice cultivation has increased by nearly 40%, while the average yield in Africa has remained stagnant (Yuan et al. 2024). Africa is partially self-sufficient in rice production with as much as 60% of the rice consumed being produced on the continent. However, the demand is increasing. Agronomic management practices in African rice cultivation, such as limiting nutrient supply and biotic factors such as diseases and pests, restrict the average rice yield to 2.4 tons ha^-1^, substantially lower when compared to Southern Asia (4.3 tons ha^-1^)(<u>FAOSTAT</u>) and far below the yield potential. Yield losses to bacterial blight (BB) of rice caused by the gram-negative bacterium, *Xanthomonas oryzae* pv. *oryzae* (*Xoo*) ranges from 4.7% in sub-Saharan Africa (Savary 2006). Increased demand and the need to improve food security demand broad-spectrum and durable resistance in local varieties. Notably, 80% of African farmers have farms less than 1 ha in size and are therefore particularly vulnerable to BB infections in their rice crop.

BB was first reported in Africa in the 1970s, and spread especially in irrigated and low-land rice growing areas in several West- and Central-African countries (Buddenhagen et al. 1979; Reckhaus 1983; Jones et al. 1991; Verdier et al. 2012), yet BB was not considered a major concern in countries including Madagascar, Tanzania, and Kenya (Schepler-Luu et al. 2023; Raveloson et al. 2023; Rabekijana et al. 2026). In 2019, two independent outbreaks in Tanzania (*Xoo* strains isolated were named iTz for introduced into Tanzania) and in Madagascar (iMg) (Schepler-Luu et al. 2023; Raveloson et al. 2023; Rabekijana et al. 2026). While detailed quantitative data are not available yet, yield losses have been estimated up to more than 20% (Ibrahim Hashim, TARI, Tanzania, pers. comm.). Therefore, there is an urgent need to prevent the spread of the disease in East Africa from the emerging *Xoo* populations. Gene deployment is the most economical and, to date, most broadly accepted approach to managing diseases, including BB, and protecting the harvest of small-scale food producers. Over 50 BB resistance genes have been identified so far, and several, including *xa13* and *Xa21* have been routinely deployed in Asian rice cultivars (Joseph et al. 2004; Sundaram et al. 2008; Ellur et al. 2016; Arra et al. 2018a, b; Sinha et al. 2023; Gautam et al. 2023). Since using individual R-genes risk disease resistance break down due to evolving *Xoo* strains, breeders pyramid combinations of resistance genes, based on the effectiveness of the R-genes against the prevalent *Xoo* strains. Combinations of three to five genes, e.g., *Xa4*, *xa5*, *Xa7*, *xa13* (*SWEET11a*), *Xa21*, *Xa22, Xa23*, *Xa33,* and *Xa38* have been used in inbred and hybrid rice cultivation in India (Sundaram et al. 2008; Gopalakrishnan et al. 2008; Basaavaraj et al. 2010; Huang et al. 2012; Balachiranjeevi et al. 2015; Arra et al. 2018b; Jamaloddin et al. 2020).

Genome editing has been used for improving rice performance, including conferring broad-spectrum BB resistance and yield improvement (Li et al. 2016; Eom et al. 2019; Oliva et al. 2019; Schepler-Luu et al. 2023). *Xoo* secretes transcription activation-like effectors (TALe) via type three secretion system (T3SS) to induce the expression of one or several clade III SWEET sugar transporters, namely *Xa13*/Os8N3/*SWEET11a*, *Xa25*/*SWEET13,* and *Xa41t*, *11N3*/*SWEET14,* which are essential for BB disease progression (Chen et al. 2010, 2012; Eom et al. 2019; Oliva et al. 2019). Editing of effector binding elements (EBEs) to prevent TALe binding in the promoters of the three SWEET susceptibility genes successfully provides rice lines with broad-spectrum resistance to BB without affecting agronomic penalties (Eom et al. 2019; Oliva et al. 2019; Schepler-Luu et al. 2023; Loo et al. 2025; Huguet-Tapia et al. 2025). Transgene-free genome-edited rice lines (SDN-1) can be introduced and deployed in countries with biosafety regulations (Buchholzer and Frommer 2023a; Schepler-Luu et al. 2023; Loo et al. 2025; Huguet-Tapia et al. 2025). However, countries such as Tanzania, Madagascar, and most neighboring countries have yet to establish biosafety regulations to allow the use of the genome-edited lines, and thus require alternative solutions to BB resistance.

Komboka (IRO5N-221) is a high-yielding, semi-aromatic rice variety developed by Kenya Agricultural & Livestock Research Organization (KALRO) (<u>BBSRC Varieties Kenya.pdf</u>) and International Rice Research Institute (IRRI, Philippines), and released in East African countries (e.g., Tanzania, Kenya, Uganda, and Burundi). Komboka possesses a yield potential of ∼7 tons ha^-1^ (Kitilu et al. 2019) with intermediate plant height, adapted to both rain-fed lowland and irrigated ecosystems (Ng’endo et al. 2022). Komboka has been described as being moderately resistant to rice blast, BB, and rice yellow mottle virus, while showing tolerance to submergence, salinity, and drought (Mutiga et al. 2021; Akwero et al. 2022; Ng’endo et al. 2022; Loo et al. 2024). However, Komboka was shown to be susceptible to iTz and Asian *Xoo* strains (Schepler-Luu et al. 2023; Loo et al. 2025). FARO-44 is a popular Nigerian rice semidwarf, early maturing (90-110 days) cultivar suited for lowland cultivation developed from crossing IR64 and Svay Rieng 64, owing to high yield potential, tolerance to drought and flood, grain quality, and cooking attributes (Amadu and Sandi-Gahun Jr 2023) (https://nifst.org/rice-cultivar-faro-44-improving-rice-productivity-in-nigeria/). NERICA-4 is an interspecific hybrid between *Oryza sativa* and *O. glaberrima*. NERICA-4 has excellent tillering ability, growth vigor, higher yield, grain, and cooking quality attributes (Jones et al. 1997; Daniel 2009). NERICA-4 has been described as susceptible to major rice diseases, including BB, although details regarding the strains tested are not provided (Loo et al. 2024). It is a semi-aromatic short-duration rice cultivar that matures in 100 days. NERICA-4 is tolerant to drought and phosphorus deficiency and is the most widely adopted upland variety, grown in more than 10 SSA countries (https://www.africarice.org/nerica).

TALe analysis of iTz and iMg strains combined with the susceptibility patterns of rice lines infected with the iTz and iMg strains indicated the presence of pthXo1-like TALe (Schepler-Luu et al. 2023; Raveloson et al. 2023; Rabekijana et al. 2026). We thus hypothesized that introgression of *xa13* (SWEET11a promoter variant) could confer resistance against iTz and iMg strains to rice varieties grown in Africa (Chu et al. 2006; Yang et al. 2006). Since Xa21 is also effective against a wide range of Asian *Xoo* strains, we deemed that introgression of *Xa21* could provide an additional layer of protection (Park et al. 2010). In the present study, elite rice breeding lines were developed in the genetic background of three popular rice varieties in Africa, i.e., Komboka, FARO-44, and NERICA-4, through marker-assisted pyramiding of four genes conferring resistance to BB, namely, *Xa1*, *Xa4*, *xa13,* and *Xa21*. Improved MABB-derived lines of Komboka, FARO-44, and NE-RICA-4 breeding lines carrying BB resistance gene(s) in different combinations, i.e., *Xa1*+*Xa4*+*xa13*+*Xa21*, *Xa1*+*Xa4*+*xa13*, *Xa1*+*Xa4*+*Xa21*, and *Xa1*+*xa13*+*Xa21* were developed. These improved lines were validated for their resistance against endemic West African *Xoo* strains as well as the strains introduced into Madagascar and Tanzania. The advanced lines are being tested in Tanzania for agronomic performance and resistance to BB, for deployment in the management of BB in rice in Africa.

## Results

### Molecular typing and disease assay indicated elite rice lines are susceptible to *Xoo* strain carrying major TALe

Komboka carries *Xa1* and *Xa4.* However, the *Xa* gene profiles in FARO-44 and NERICA-4 were not known. We checked for the presence of select *Xa* genes, i.e., *Xa1, Xa4, xa13,* and *Xa21* in all three rice lines using gene-linked markers. The primer pairs XaL/F1R1 produce a 270 bp amplicon indicative of the presence of *Xa1* gene (Ji et al. 2020) whereas MP1/MP2 primer pair produce a 180 bp amplicon indicative of the presence of *Xa4* gene (Sun et al. 2003). The Kitaake rice variety which does not carry both *Xa1* and *Xa4* was used as a negative control (Figure 1). The XaL/F1R1 pair amplified two amplicons between 200 and 300 bp in Komboka and FARO-44 but only a 1000 bp amplicon in Kitaake and NERICA-4 (Figure 1a). The MP1/MP2 markers amplified an amplicon smaller than 200 bp in Komboka and NERICA-4, whereas a smaller amplicon between 75 and 200 bp in Kitaake and FARO-44 (Figure 1b). Molecular typing of *Xa* genes confirmed that the resistance alleles *Xa1* and *Xa4* genes are present in Komboka, *Xa1* is present in FARO-44, and *Xa4* in NERICA-4. Using the same approach, we genotyped for the presence of *Xa1* and *Xa4* in other elite rice varieties, and found that FARO-66 and FARO-67 but not IR64, MTU1010, or NERICA L-19 carry *Xa1*; whereas FARO-66, FARO-67, IR64, and MTU1010, but not NERICA L-19 carry *Xa4* (Figure 1a-b).

**Figure 1.**
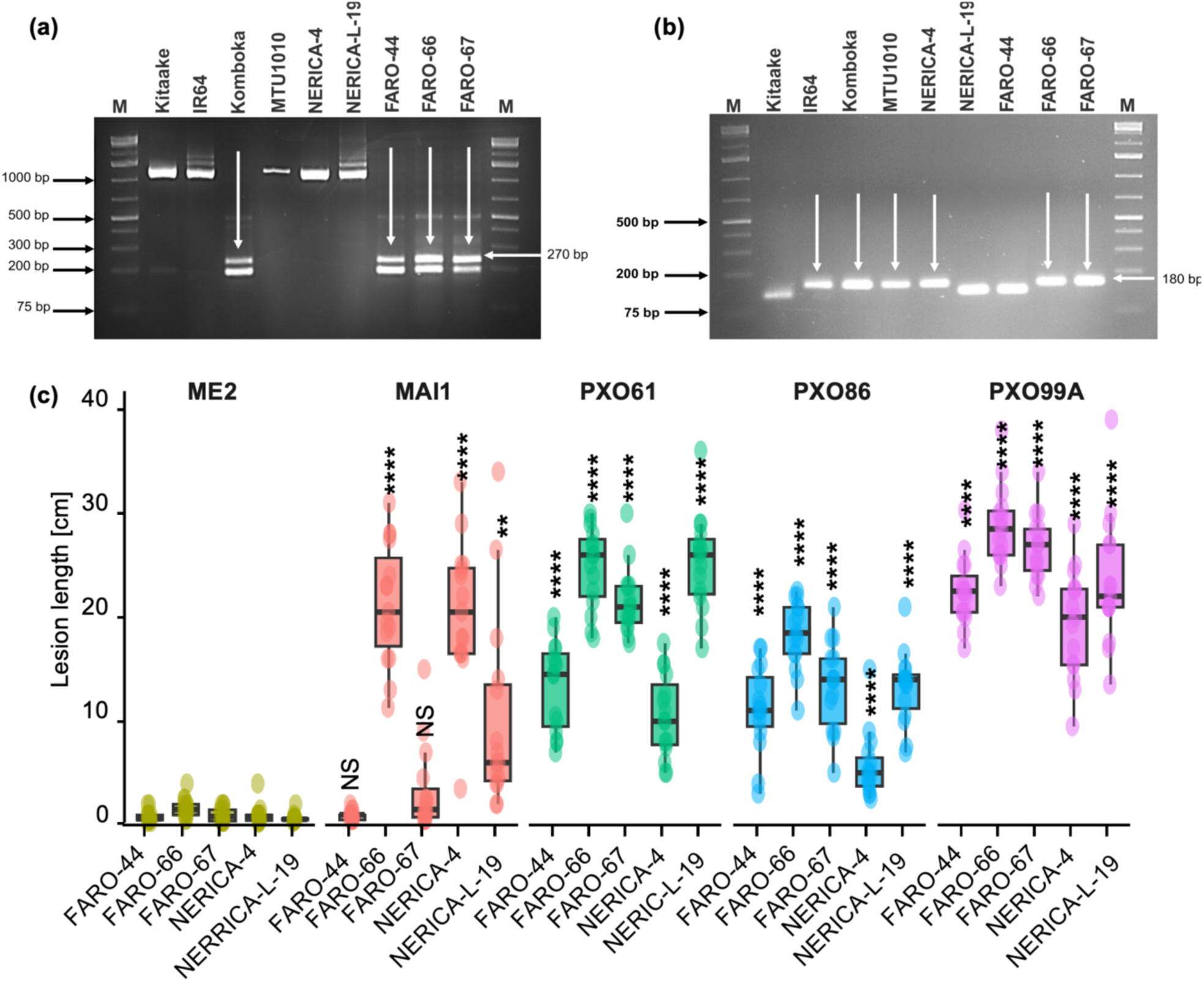
Characterization of select *Xa* gene outfit and BB susceptibility in elite rice cultivars. **a)** genotyping of *Xa1* resistance gene; **b)** genotyping of *Xa4* resistance gene; **c)** Virulence of Asian *Xoo* strains on popular African elite rice varieties NERICA-4, NERICA-L-19, FARO-44, FARO-66, and FARO-67 using the Kauffman clipping assay. P-values were calculated using Student’s T-test against ME2 of the respective cultivar. NS-not significant, ** - 0.01, **** - 0.0001. N > 10, representative dataset from three independent repeats with comparable results (see Figure S1).

In addition to genotyping, the five African rice cultivars NERICA-4, NERICA-L-19, FARO-44, FARO-66 and FARO-67 were screened for BB susceptibility using four representative virulent *Xoo* strains from Asia and Africa carrying major TALe known to induce *SWEET11a*, *SWEET13,* and *SWEET14*, i.e., PXO99^A^ (*SWEET11a-*inducing PthXo1), PXO61 (*SWEET13* (japonica cultivar)-inducing PthXo2, *SWEET14-*inducing PthXo3), PXO86 (*SWEET14-*inducing AvrXa7), and MAI1 (*SWEET14-*inducing TalF); and an avirulent strain PXO99A[ME2] as a negative control. All rice cultivars tested were susceptible to moderately susceptible towards all virulent *Xoo* strains tested except for FARO-44 and FARO-67, which were resistant to MAI1, and NERICA-4 which was moderately resistant to PXO86 (Figure 1c, Figure S1). The pathogen assay indicated that the rice lines were still susceptible to *Xoo* carrying major TALe despite carrying both *Xa1* and *Xa4*, suggesting a need for pyramiding additional *Xa* genes.

### African elite rice varieties carry TALe binding EBEs on *SWEET11a, 13*, and *14* promoters

Since modifications of the EBE were effective against TALe-binding required for BB disease progression, we analyzed the promoters of *SWEET11a, SWEET13,* and *SWEET14* in the rice varieties for potential endogenous EBE variants. The EBEs for PthXo1 in *SWEET11a,* and TalC, PthXo3, AvrXa7, and TalF in *SWEET14* were identical in all six rice cultivars. NERICA-4 carried an 11 bp deletion and T/C substitution in the EBE for PthXo2 in *SWEET13* (Figure 2). Our analysis thus indicated that the resistance of FARO-44 and FARO-67 to MAII, and resistance of NERICA-4 to PXO86 were not consequences of modification in EBEs upstream of *SWEET14,* rather possibly a phenotype conferred by yet unknown endogenous resistance genes.

**Figure 2:**
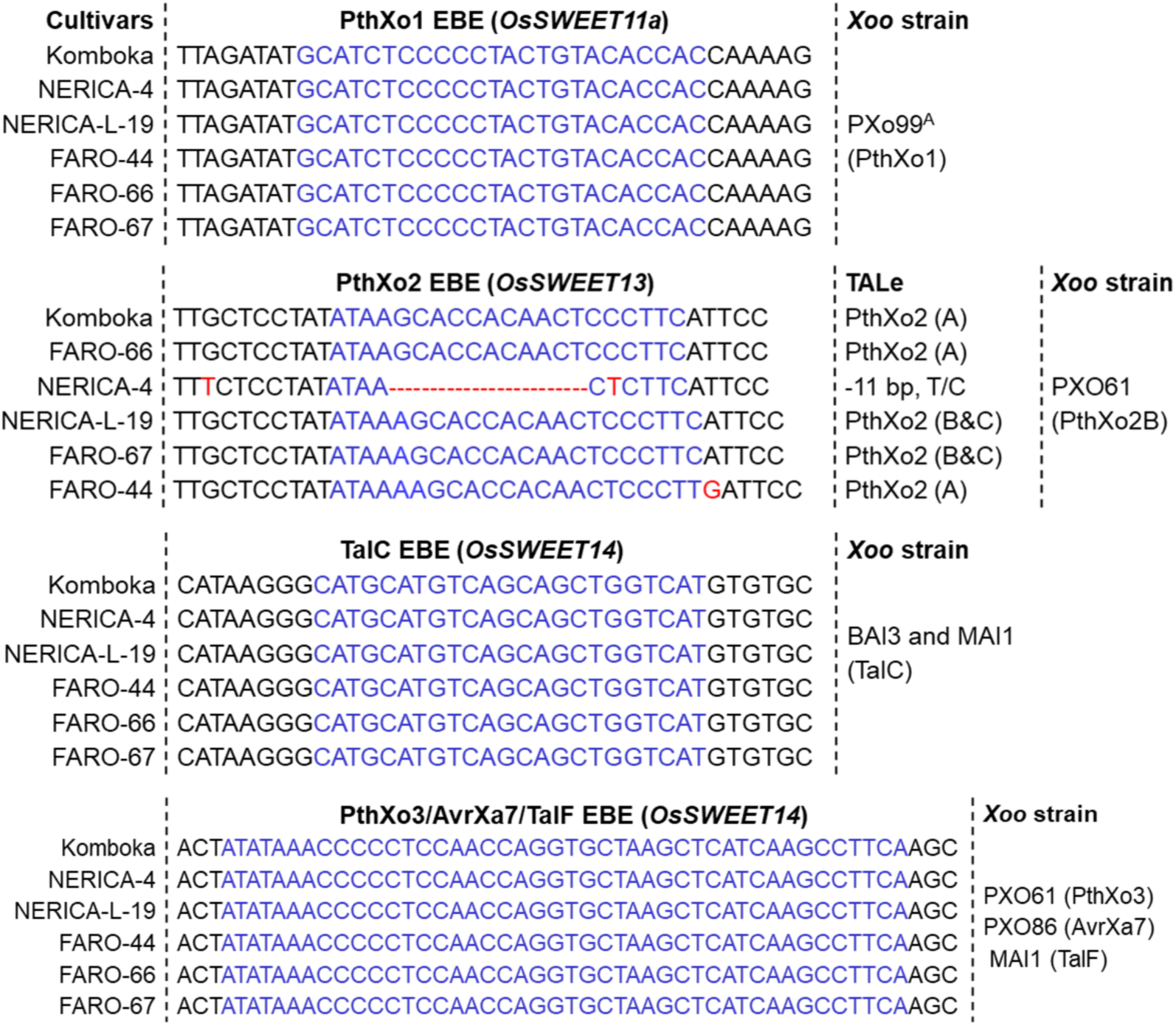
Major TALe effector-binding elements (EBEs) in the SWEET promoters of select African rice cultivars. EBE sequences are highlighted in blue; single-nucleotide polymorphisms are highlighted in red.

### Introgression of *xa13* and *Xa21* resistance genes into Komboka via MABB using IRBB55

Since TALeome analysis of iTz and iMg strains indicated the presence of PthXo1-like TALe which likely largely promoted susceptibility in the tested rice varieties (Figure 1c), we hypothesized that introgression of *xa13* and *Xa21* could confer Komboka resistance against the introduced strains (Huang et al. 1997; Sundaram et al. 2008; Arra et al. 2017; Schepler-Luu et al. 2023; Raveloson et al. 2023; Rabekijana et al. 2026). IRBB55, which carries *xa13* and *Xa21* (Joseph et al. 2004), was crossed with Komboka to introgress *xa13* and *Xa21* into Komboka. The resulting F_1_ plants obtained (Figure 3) were genotyped for the presence of *xa13* and *Xa21* using established markers, i.e., Xa13-prom for SWEET11a/*xa13* (Hajira et al. 2016) and pTA248 for *Xa21* (Ronald et al. 1992) (Figure 3, Table S1). Amplicon sizes amplified using individual gene-specific markers in the donor parent (IRBB55) and the recipient parent (Komboka) were used to benchmark hetero- or homozygosity in the F_1_ progeny. The Xa13-prom marker is co-dominant and produced an amplicon with a length matching the expected 273 bp in Komboka, and 488 bp in IRBB55 (Figure 4).

**Figure 3:**
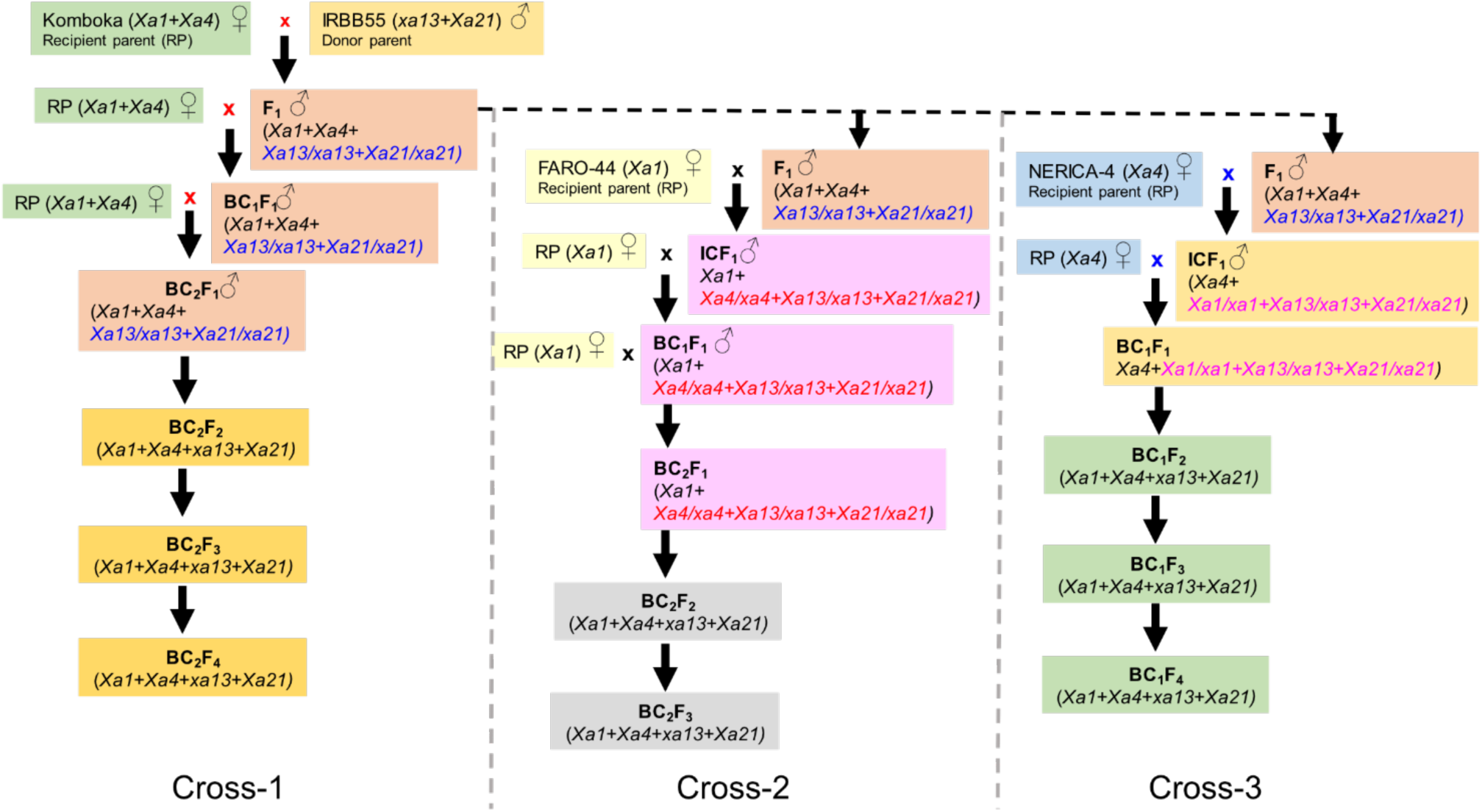
A Rice crossing scheme adopted in this study for the development of broad-spectrum BB resistant Komboka, FARO-44 and NERICA-4 lines. Cross 1: Two major BB resistance genes *xa13* and *Xa21* from IRBB55 were introduced into Komboka; Cross-2: Three BB resistance genes *Xa4, xa13,* and *Xa21* from Cross-1 were introgressed into FARO-44; Cross-3: Three BB resistance genes *Xa1*, *xa13* and *Xa21* from Cross-1 were introgressed into NERICA-4. The BB R-genes were introgressed into Komboka, FARO-44, and NERICA-4 are highlighted with blue, red and purple, respectively.

**Figure 4:**
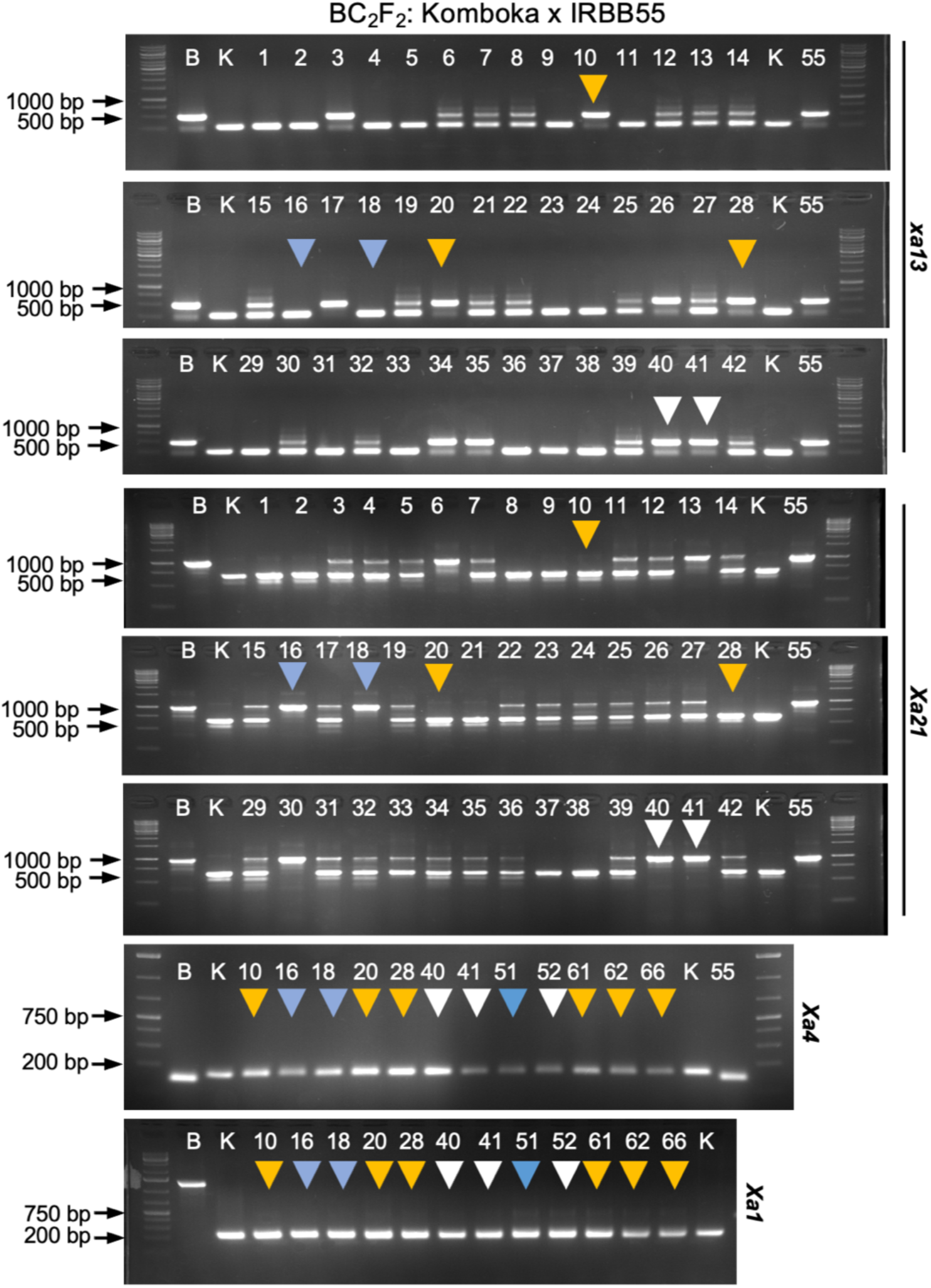
Molecular typing of *Xa* genes in BC_2_F_2_ progenies derived from IRBB55 x Komboka MABB. Genotyping for indicated *Xa* genes in BC_2_F_2_ progenies derived from BC_1_F_2_ line 5 (representative line) using XaL (*Xa1*), MP1/MP2 (Xa4), xa13prom (*xa13*), and pTA248 (*Xa21*) markers (Table S1). White arrow heads indicate lines carrying *Xa1+Xa4+xa13 Xa21*, yellow arrow heads indicate lines carrying *Xa1+Xa4+ xa13,* blue arrow heads indicate lines carrying *Xa1+Xa4+ Xa21.* Numbers above each lane represent each BC_2_F_2_ progeny line, K-Komboka, B-IRBB55.

The F_1_ progenies were advanced and backcrossed to BC_2_F_1_ (Figure 3). A total of 11 F_1_ plants were advanced, of which six were heterozygous for both *xa13* and *Xa21*. One of the F_1_ plants was used as a donor and back-crossed with the recurrent parent (Komboka) to generate BC_1_F_1_. Out of 22 BC_1_F_1_ plants analyzed for the presence of *xa13* and *Xa21*, three plants were heterozygous for *xa13* and *Xa21*. One of the BC_1_F_1_ plants was used as a donor and backcrossed with the Komboka to generate BC_2_F_1_ seeds. Three BC_2_F_1_ lines (i.e., lines 5, 8 & 12, Figure S2a) were heterozygous for *xa13* and *Xa21* as identified through foreground selection and thus, advanced to BC_2_F_2_. Since Komboka carries endogenous *Xa1* and *Xa4*, the retention of *Xa1* and *Xa4* and the addition of *xa13* and *Xa21* were tested in BC_2_F_2_ (Figure 4a). A total of 208 BC_2_F_2_ plants from three progenies were screened for the presence of the two introgressed genes *xa13*, and *Xa21,* and the endogenous *Xa1* and *Xa4* using XaL-F1R1 for *Xa1* (Ji et al. 2020) and MP1/MP2 for *Xa4* (Sun et al. 2003) (Table S1). Two plants out of 66 BC_2_F_2_ derived from line 5, five plants of 89 BC_2_F_2_ derived from line 8, and one plant of 53 BC_2_F_2_ derived from line 12 were homozygous for all four *Xa* genes (Figure 4; Table 1). Besides homozygous plants carrying all four *Xa* genes, homozygous plants with three gene combinations were also identified (Figure 4a, Table 1). Individual BC_2_F_2_ homozygous plants with three or four *Xa* gene combinations were advanced by self-pollination to generate BC_2_F_3_.

**Table 1.** List of improved breeding lines in the backgrounds of the elite rice varieties Komboka, FARO-44 and NERICA-4 developed by MABB.

| Designation | R-gene combination | Backcross generation | Plant height [cm]<br>mean±SE | Days to seed maturity<br>mean±SE | Grain type | RPGR (%) |
| --- | --- | --- | --- | --- | --- | --- |
| **Komboka improved breeding lines** |  |  |  |  |  |  |
| 5-66 | *Xa1Xa1+Xa4Xa4+xa13xa13* | BC<sub>2</sub>F<sub>3</sub> | 104.4 ± 0.6 | 146.0 ± 1.6 | MS | 94.2 |
| 8-3 | *Xa1Xa1+Xa4Xa4+xa13xa13* | BC<sub>2</sub>F<sub>3</sub> | 115.3 ± 1.6 | 147.3 ± 0.6 | MS | 91.3 |
| 5-52 | *Xa1Xa1+Xa4Xa4+xa13xa13+Xa21Xa21* | BC<sub>2</sub>F<sub>3</sub> | 124.2 ± 1.2 | 145.8 ± 1.7 | MS | 91.3 |
| 8-32 | *Xa1Xa1+Xa4Xa4+xa13xa13+Xa21Xa21* | BC<sub>2</sub>F<sub>3</sub> | 121.0 ± 0.6 | 146.8 ± 0.9 | MS | 88.4 |
| 8-65 | *Xa1Xa1+Xa4Xa4+xa13xa13+Xa21Xa21* | BC<sub>2</sub>F<sub>3</sub> | 114.3 ± 1.3 | 146.5 ± 0.6 | MS | 94.2 |
| 12-15 | *Xa1Xa1+Xa4Xa4+xa13xa13+Xa21Xa21* | BC<sub>2</sub>F<sub>3</sub> | 130.1 ± 0.5 | 135.8 ± 0.9 | MS | 88.4 |
| Komboka | *Xa1Xa1+Xa4Xa4+Xa13Xa13+xa21xa21* | cultivar | 124.5 ± 3.5 | 149.0 ± 1.6 | MS |  |
| IRBB 55 | *xa1xa1+xa4xa4+xa13xa13+Xa21Xa21* | Isogenic IR24 line | 122.8 ± 2.6 | 140.5 ± 0.6 | MS |  |
| **FARO-44 improved breeding lines** |  |  |  |  |  |  |
| 31-15 | *Xa1Xa1+Xa4Xa4+xa13xa13* | BC<sub>2</sub>F<sub>3</sub> | 123.0 ± 0.9 | 133.5 ± 0.6 | LS |  |
| 19-70 | *XaXa1+xa13xa13+Xa21Xa21* | BC<sub>2</sub>F<sub>3</sub> | 123.7 ± 0.9 | 135.0 ± 1.9 | LS |  |
| 19-80 | *Xa1Xa1+Xa4Xa4+xa13xa13+Xa21Xa21* | BC<sub>2</sub>F<sub>3</sub> | 124.2 ± 0.5 | 132.8 ± 0.9 | LS |  |
| FARO 44 | *Xa1Xa1+xa4xa4+Xa13Xa13+xa21xa21* | cultivar | 122.9 ± 1.1 | 134.3 ± 1.4 |  |  |
| **NERICA-4 improved breeding lines** |  |  |  |  |  |  |
| 13-4 | *Xa1Xa1+Xa4Xa4+Xa21Xa21* | BC<sub>1</sub>F<sub>4</sub> | 141.8± 1.2 | 129.8 ± 0.9 | LS |  |
| 13-11 | *Xa1Xa1+Xa4Xa4+xa13xa13* | BC<sub>1</sub>F<sub>4</sub> | 140.9± 0.8 | 131.3 ± 1.0 | LS |  |
| 4-49 | *Xa1Xa1+xa13xa13+Xa21Xa21* | BC<sub>1</sub>F<sub>4</sub> | 142.1± 1.1 | 130.5 ± 0.9 | LS |  |
| 13-34 | *Xa1Xa1+Xa4Xa4+xa13xa13+Xa21Xa21* | BC<sub>1</sub>F<sub>4</sub> | 141.0± 0.6 | 132.0 ± 0.9 | LS |  |
| NERICA-4 | *xa1xa1+Xa4Xa4+Xa13Xa13+xa21xa21* | cultivar | 143.1± 1.3 | 131.0 ± 1.1 | LS |  |

**MS:** Medium slender; **LS:** Long slender; **RP:** Recipient parent; **DP:** Donor parent; **RPGR:** Recurrent parent genome recovery.

Linkage drag is one of the critical factors in breeding that restricts the fitness of developed breeding lines of the recipient cultivar. MABB breeding allows for monitoring the target genes from a donor parent chromosome segment during each backcross through polymorphic simple sequence repeat (SSR) molecular markers. To determine parent genome recovery (RPG), screening of polymorphic SSRs was performed using previously established SSR markers (Figure 5a). The linkage drag for Komboka BC_2_F_3_ homozygous breeding lines was delimited to 0.2 Mb for *xa13* (each 0.1 Mb at upstream and downstream region) and ∼ 0.4 Mb for *Xa21* (0.1 Mb at the upstream and 0.3 Mb at the downstream region) (Figures 5b, c). Eighty-five polymorphic SSR markers for Komboka, 43 polymorphic SSR markers for FARO-44, and 74 polymorphic SSR markers for NERICA-4 distributed across the 12 rice chromosomes (i.e., ∼2–19 SSRs) were identified (Table S2-S4). In sum, the background screen indicated that the homozygous BC_2_F_3_ plants had an RPG with a range of 88.4-94.2%, and exhibited phenotypic similarity to Komboka (Figure 5; Table 1, Figure S3).

**Figure 5:**
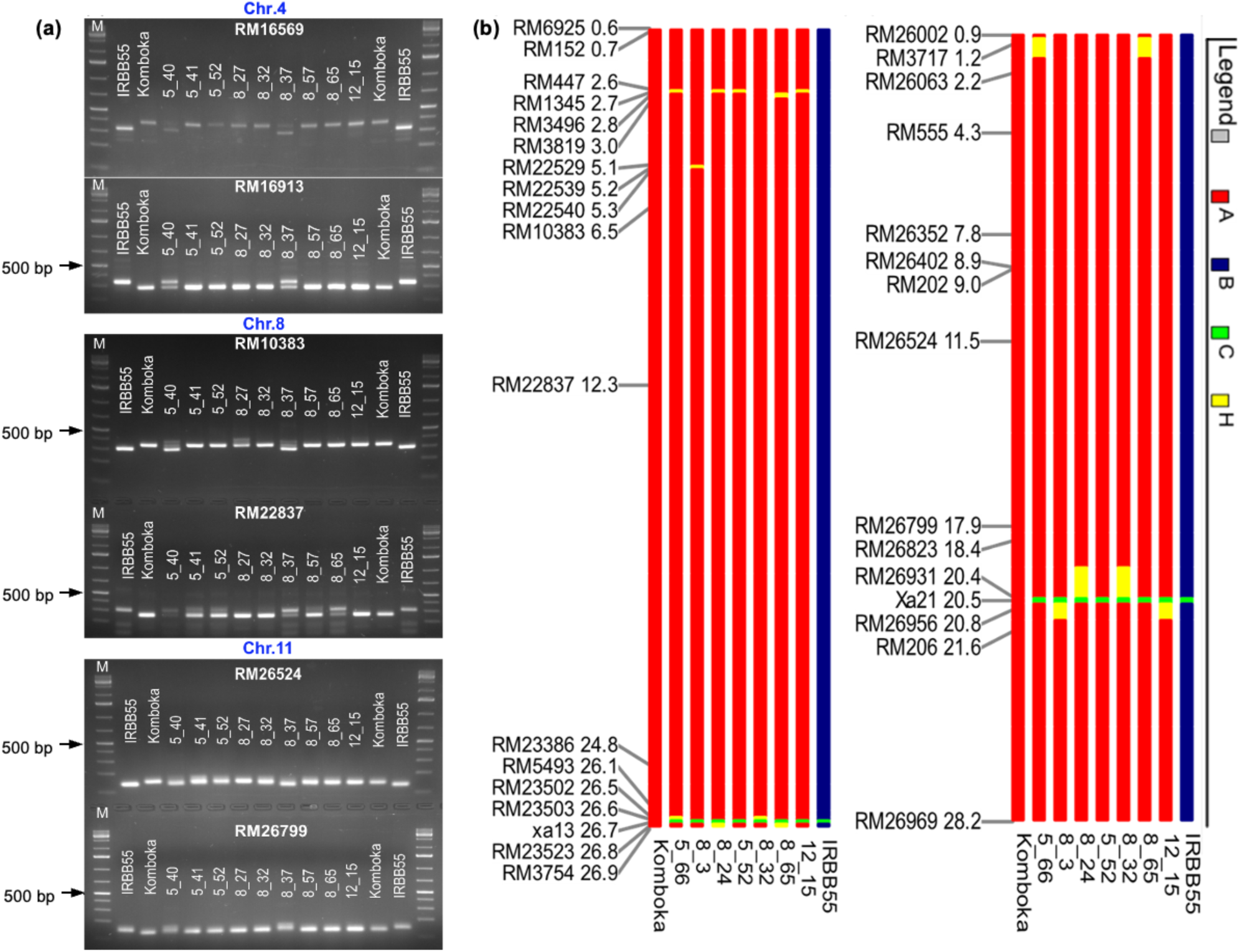
Linkage drag associated with *xa13* and *Xa21* in the genetic background of BC_2_F_3_ IRBB55 x Komboka MABB-derived lines. **(a)** Representative gel images for background selection with polymorphic SSR markers for recurrent parent genome recovery. **(b)** Analysis of genome introgression associated with the *xa13*-genomic region limited to ∼0.5 Mb introgressed from IRBB55 in the selected MABB lines. Analysis of genome introgression associated with the *Xa21*-genomic region limited to ∼0.7 Mb introgressed from IRBB55 in the selected MABB lines. The position of each polymorphic SSR marker in Mb on chromosomes 8 and 11. In legend, A: Komboka; B: IRBB55; C: gene location; H: heterozygous.

### Introgression of *Xa* genes into FARO-44 and NERICA-4 via MABB using IRBB55/Komboka

Since NERICA-4 carries a 11 bp deletion in the EBE of PthXo2, and thus will be resistant to *Xoo* strains carrying PthXo2 TALe, as well as FARO-44 were chosen for introgression with *xa13*, *Xa21,* and *Xa1* or *Xa4* (Figure 3). To reduce the number of crosses required to introgress three *Xa* genes into FARO-44 and NERICA-4, we used an F_1_ progeny from the IRBB55 x Komboka cross carrying all four *Xa* genes (Figure 3, 4) as the donor parent. Two out of 18 F_1_ progenies derived from the FARO-44 x IRBB55/Komboka (i.e., lines 3 and 15, Figure S2b) were heterozygous for *xa13* and *Xa21* as indicated by successful amplification of amplicons as expected sizes using the MP1/MP2 Xa13-prom and pTA248 primers (Ronald et al. 1992; Sun et al. 2003; Hajira et al. 2016). The F_1_ progeny line 3 was used as a donor and backcrossed with FARO-44 to obtain BC_1_F_1_. Two resulting plants (i.e., lines 11 and 15) out of 18 BC_1_F_1_ plants were heterozygous for *Xa4*, *xa13*, *Xa21*. The BC_1_F_1_ plant line 11 was subsequently used for backcrossing with FARO-44 to obtain BC_2_F_1_. Two plants (i.e., lines 19 and 31) out of 38 BC_2_F_1_ plants were heterozygous for *Xa4*, *xa13*, and *Xa21* and were advanced to obtain BC_2_F_2_ generation by self-pollination. A total of 96 plants from each BC_2_F_2_ progeny line 19 and 31 were screened. One plant from BC_2_F_2_ line 19, named as line 19-80 was homozygous for all four *Xa* genes (Figure 6a). Plant 19-80 was advanced to generate BC_2_F_3_ seeds. BC_2_F_2_ plants with three *Xa* gene combinations, *Xa1*+*xa13*+*Xa21* and *Xa1*+*Xa4*+*xa13* were also selected as potential candidates for broad-spectrum BB resistance (Table 1).

**Figure 6:**
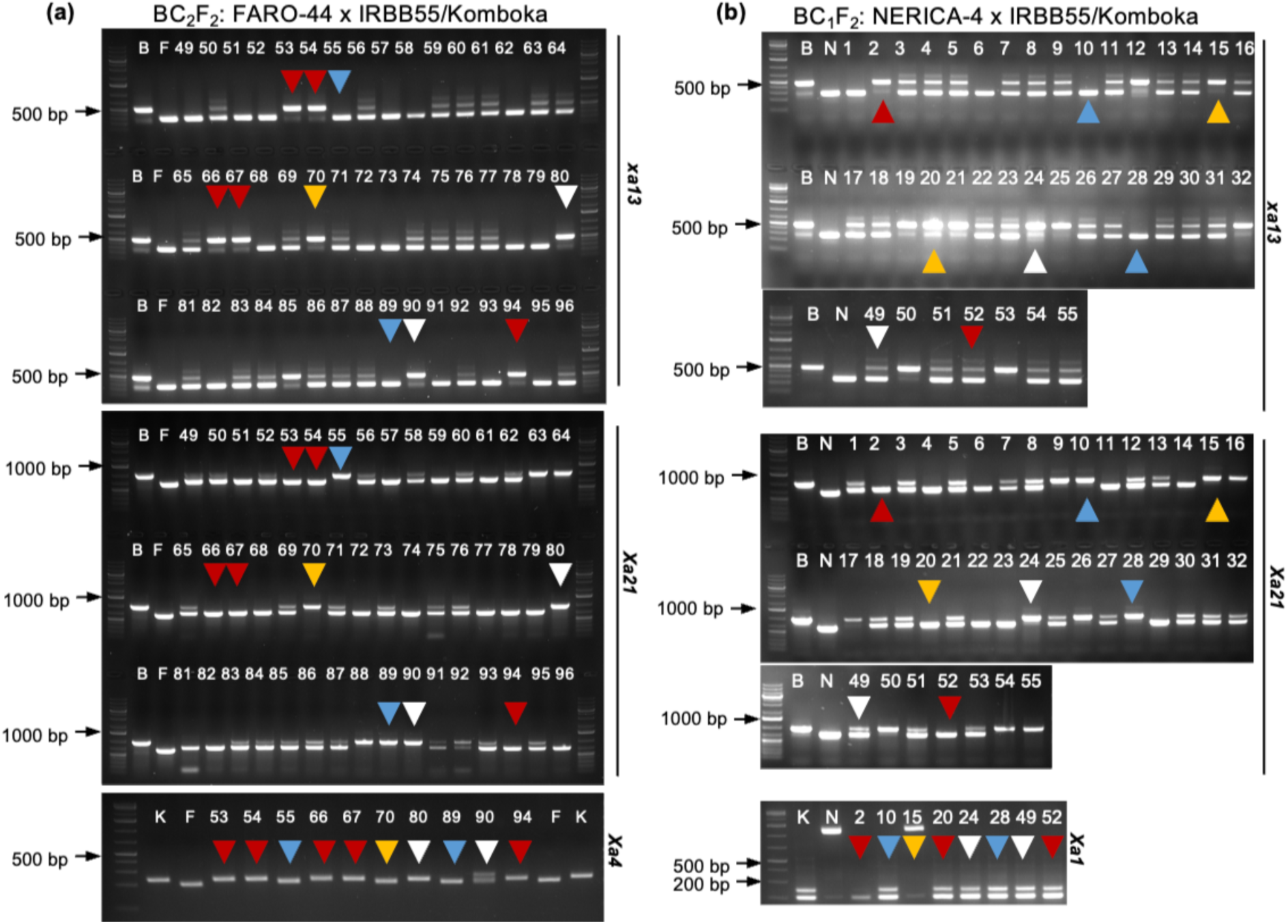
Molecular typing of *Xa* genes in progenies derived from FARO-44 or NERICA-4 x IRBB55/ Komboka MABB lines. Genotyping for indicated *Xa* genes in **(a)** BC_2_F_2_ progenies derived from BC_1_F_1_ line 19 of FARO-44 x IRBB55/Komboka (representative line). White arrow heads represent lines carrying all four *Xa1+Xa4+xa13+Xa21*, yellow arrow heads indicate lines carrying *Xa1+ xa13+Xa21,* blue arrow heads indi-cate lines carrying *Xa1*+*Xa21,* red arrow heads indicate lines carrying *Xa1+Xa4+ xa13;* in **(b)** BC_1_F_2_ progenies derived from BC_1_F_1_ line 4 of NERICA-4 x IRBB55/Komboka (representative line). White arrow heads indicate lines carrying all four *Xa1+Xa4+xa13+Xa21*, yellow arrow heads indicate lines carrying *Xa4+ xa13+Xa21,* blue arrow heads indicate lines carrying *Xa1*+*Xa4*+*Xa21,* red arrow heads indicate lines carrying *Xa1+Xa4+ xa13.* Numbers above each lane represent each BC_2_F_2_ progeny line, K-Komboka, B-IRBB55.

To introgress *Xa1*, *xa13*, *Xa21* into NERICA-4, we employed the same strategy used for FARO-44 whereby an F_1_ progeny from IRBB55 x Komboka carrying all four *Xa* genes were used as donor parent (Figure 3). Four resulting F_1_ progenies (lines 2, 4, 5, and 9, Figure S2c) out of 14 progenies were identified as heterozygotes for *Xa1*, *xa13,* and *Xa21,* in foreground selection using molecular markers. The observed number of heterozygous plants is higher than the expected, likely a statistical bias due to the low number of plants analyzed. Since NERICA-4 is derived from an interspecific hybridization between *Oryza sativa* x *Oryza glaberrima*, the F_1_ plants derived from NERICA-4 x IRBB55/Komboka were evaluated for seed-setting and had a rate of more than 50% (Figure S4), indicating high fertility for subsequent crossing. Among the 24 BC_1_F_1_ plants, five plants (lines 4, 9, 13, 19, and 23) were heterozygous for *Xa1*, *xa13,* and *Xa21*. These five selected plants were advanced by self-pollination to obtain BC_1_F_2_. Two BC_1_F_2_ progenies (lines 4 and 13) were chosen as subsequent donor parents based on the plant phenotype, percentage of seed setting (>50%) and homozygous for *Xa1*, *Xa4*, *xa13*, and *Xa21*. Two plants (lines 4-24, and 4-52) out of 57 from BC_1_F_2_ progenies derived from line 4, and one plant (line 13-34) out of 37 BC_1_F_2_ progenies derived from line 13, were homozygous for *Xa1*, *Xa4*, *xa13,* and *Xa21* (Figure 6b) thus, advanced to obtain BC_1_F_4_. Agronomic and morphological characteristics were evaluated from individual progenies of BC_1_F_2_ and, based on performance, promising breeding lines were chosen for further evaluation.

### MABB-derived improved Komboka, FARO-44, and NERICA-4 lines are resistant to iTz and iMg *Xoo* strains

The improved BC_2_F_3_ backcross breeding lines of Komboka were evaluated for disease resistance with *Xoo* strains collected from Tanzania (iTz) and Madagascar (iMg). A total of six BC_2_F_3_ breeding lines, which possess three gene combinations, *Xa1*+*Xa4*+*xa13* (5-66, and 8-3), and four gene combinations, *Xa1*+*Xa4*+*xa13*+*Xa21* (lines 5-52, 8-32, 8-65, and 12-15) were tested against 14 iTz and two iMg strains, and PXO86 (AvrXa7) previously shown virulent on Komboka (Schepler-Luu et al. 2023) was used as a positive control. All improved BC_2_F_3_ breeding lines of Komboka have higher resistance levels to the tested *Xoo* strains, especially four gene combinations (lines 5-52, 8-32, 8-65, and 12-15) indicated by the significantly reduced lesion length compared to the parental lines (Figures 7a, Figure S5-S7). Two BC_2_F_3_ breeding lines (lines 5-66, and 8-3) with three gene combinations (*Xa1*+*Xa4*+*xa13*) were susceptible to the PXO86. Four BC_2_F_3_ breeding lines with four gene combinations (*Xa1*+*Xa4*+*xa13+Xa21*) were moderately resistant to PXO86.

**Figure 7.**
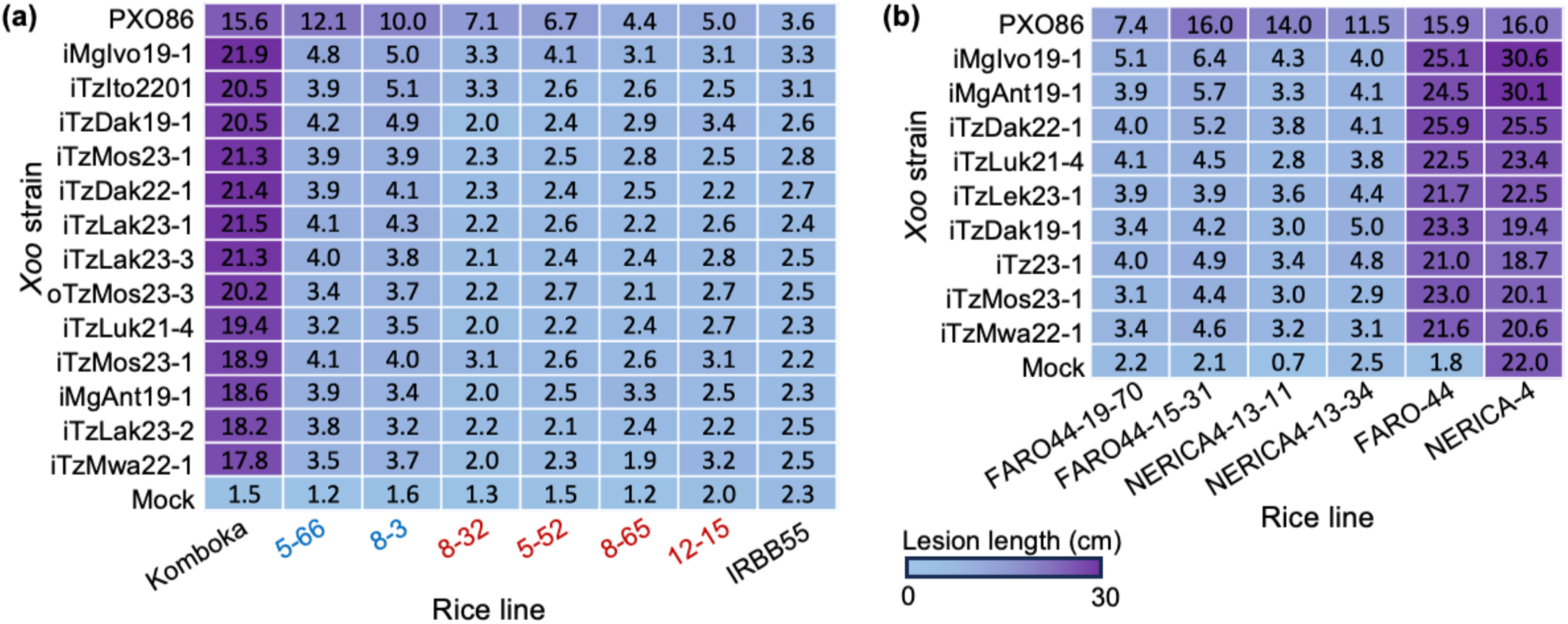
Virulence profiles of *Xanthomonas oryzae* pv. *oryzae* isolates from Tanzania (iTz) and Madagascar (iMg) on. **(a)** Komboka x IRBB55; and **(b)** FARO-44 or NERICA-4 x IRBB55/Komboka MABB-derived lines with introgressed *Xa* genes. The heatmap displays mean lesion length (in cm) for each plant line clip-infected with the indicated *Xoo* strains. Mean lesion length calculated from three independent experiments, N> 8 in each repeat. Lesion length was measured 21 days post-infection (see Figure S6 for statistical analysis). In **(a)**, lines indicated in blue carry a *Xa1+Xa4+xa13*, and lines indicated in red carry *Xa1+Xa4+xa13*+*Xa21*.

Similarly, the BC_2_F_3_ offsprings from FARO-44 x IRBB55/Komboka and BC_1_F_4_ plants from NERICA-4 x IRBB55/Komboka were evaluated for BB disease resistance with ten iTz, two iMg strains and PXO86 (Figure 7b). The FARO-44 BC_2_F_3_ homozygous backcross lines 19-70 (*Xa1*+*xa13*+*Xa21*) and 15-31 (*Xa1*+*Xa4*+*xa13*), and NERICA-4 BC_1_F_4_ homozygous backcross lines 13-11 (*Xa1*+*Xa4*+*xa13*), and 13-34 (*Xa1*+*Xa4*+*xa13*+*Xa21*) were resistant to seven iTz strains and two iMg strains, respectively (Figure 7b). Similar to their parental lines, FARO-44 backcross lines 15-31 (*Xa1*+*Xa4*+*xa13*) and 13-11 (*Xa1*+*Xa4*+*xa13*), and NERICA-4 backcross line 13-34 (*Xa1*+*Xa4*+*xa13*+*Xa21*) were moderately susceptible to PXO86, except for the three gene combination FARO-44 line 19-70 (*Xa1*+*xa13*+*Xa21*) which showed moderate resistance. These differences may be explained by the role of the genetic background for the resistance, e.g., the three gene combination *xa5*+*xa13*+*Xa21* in Improved Samba Mahsuri showed resistance to respective strains, whereas in the Triguna cultivar carrying the three R genes was more susceptible (Sundaram et al. 2009).

## Discussion

Besides high quality standards in plant pest quarantine and improved management practices, genetic resistance is an economic and effective way to prevent plant disease epidemics and spread, host plant resistance offers to introduce significant resistance genes in relevant rice cultivars (Sundaram et al. 2008; Laha et al. 2009, 2016; Jamaloddin et al. 2020; Gopalakrishnan et al. 2022; Loo et al. 2024). Editing of all known EBEs in SWEET promoters through CRISPR-Cas has provided broad-spectrum resistance to a all currently known *Xoo* races/pathotypes in the japonica variety Komboka and multiple elite cultivars (Li et al. 2012, 2025; Blanvillain-Baufumé et al. 2017; Eom et al. 2019; Oliva et al. 2019; Gupta et al. 2023, 2026; Schepler-Luu et al. 2023). However, due to lack of harmonization of genome-editing biosafety regulations, genome-edited crops cannot be cultivated in all countries (https://www.editagenome.org)(Buchholzer and Frommer 2023b). A classical approach for protecting rice from BB has been to evaluate the efficacy natural resistance gene variants against a panel of Xoo strains (Laha et al. 2009; Arra et al. 2017). Over 50 BB resistance genes have been reported (Sinha et al. 2023), and a few resistance genes are routinely introgressed in BB-susceptible rice cultivars and hybrid parental lines (Sundaram et al. 2008; Balachiranjeevi et al. 2015; Ellur et al. 2016; Arra et al. 2018a, b; Fiyaz et al. 2022). The ability to induce sucrose transporting SWEETs appears to be critical for virulence, therefore we surmise that the combination of genetic EBE variants as made possible by genome editing will be more robust since single SWEET resistance alleles can be broken by circumventing a particular SWEET through induction of another. This can be achieved for example by horizontal transfer of TALe from one strain to another. However classical breeding cannot combine all SWEET EBE variants in one line due to tight linkage of some, i.e. multiple EBEs in are used by different strains in the promoter of SWEET14 (Eom et al. 2019; Schepler-Luu et al. 2023). However, to provide a rapid fix for countries that do not have Genome editing regulations, in particular East African countries that have been impacted by outbreaks resulting from the inadvertent introduction of Asian strains, we here tried to make use of the information gained from the characterization of the strains introduced into Tanzania and Madagascar to develop elite varieties using MABB (Schepler-Luu et al. 2023; Rabekijana et al. 2026). Notably, the development a new variety through conventional breeding takes at least 7 years (Gopalakrishnan et al. 2022). Stacking multiple resistance genes through classical breeding methods is often challenging due to linkage drag, dominance, and epistatic effects (Huang et al. 2012; Gopalakrishnan et al. 2022). Marker-assisted backcross breeding (MABB) enables the introgression of desirable traits into elite rice varieties (two generations/year) within 2-3 years to generate BC_2-3_ generation homozygous breeding lines under controlled greenhouse conditions.

We here first evaluated several African elite varieties for the presence of R genes and for the susceptibility to a panel of related strains isolated from the outbreaks in Tanzania and Madagascar. Since the introduced strains carry the Asian pThXo1 TALe, we introduced *xa13*, and as an additional layer of protection we also added the widely used *Xa21* R gene through MABB in a stepwise process. We first crossed Komboka, which carries *Xa1* and *Xa4* to IRBB55, which carries both homozygous versions of *xa13* and *Xa2*1, and tracked the introgression until BC_2_F_2_ using fore- and background selection by MABB. In parallel we used the F_1_ from the Komboka x IRBB55 cross to combine all four R genes in NERICA-4 and FARO-44. NERICA-4 was backcrossed until BC_1_F_2_, and FARO-44 to BC_2_F_2_ using fore- and background selection by MABB as well. Finally, the resistance of four BC_2_F_3_ Komboka x IRBB55 lines, two three-way-cross lines in BC_1_F_4_ (F_1_ Komboka/IRBB55 x NE-RICA-4) and two three-way-cross lines in BC_2_F_4_ (F_1_ Komboka/IRBB55 x FARO-44) to a panel of ten to 14 strains (including Asian, iMg and iTz strains) using the Kaufmann clipping assay. The new lines were resistant to all strains. Subsequently, the lines were transferred to breeders in Tanzania and Madagascar for further development and registration.

In summary, the present study reports the combination of four major BB resistance genes into the Komboka, FARO-44, and NERICA-4 cultivars using MABB. Since the improved MABB breeding lines are resistant to independent isolates of the introduced iTz and iMg *Xoo* strains from East Africa, as well as to Asian *Xoo* strains (PXO99^A^ and PXO86), they may help to protect the harvest of small-scale food producers in African countries that do not yet have established regulations that would allow them to use the genome-edited broaspectrum BB resistant SWEET-promoter lines. The lines have already successfully been transferred to TARI in Tanzania and FOFIFA in Madagascar, other transfers being processed. We note that likely, the edited lines distributed to Kenia will provide broader and more robust resistance.

## Materials and Methods

### Plant material

Komboka, a recent African rice cultivar, was selected as the recipient parent. Komboka possesses two dominant BB resistances, *Xa1* and *Xa4* (Schepler-Luu et al. 2023). Recently introduced *Xoo* isolates from Tanzania and Madagascar were found to be virulent on Komboka. (Schepler-Luu et al. 2023). The area-wise cultivation of Komboka is increasing in East African countries. Therefore, it is important that resistant Komboka be developed to *Xoo* strains introduced into Africa from Asia. FARO-44 is an irrigated lowland high-yield potential rice cultivar released in sub-Saharan Africa developed by the International Rice Research Institute, Philippines (Manasseh et al. 2018). This variety has become more popular among farmers in Kenya due to its grain quality attributes. The introduction of Asian *Xoo* strains into East Africa, local rice cultivars were severely affected by BB, and the FARO-44 became highly susceptible to BB when tested against the *Xoo* strains (unpublished data). NERICA-4, an upland rice cultivar in African countries is well-adapted among NERICA cultivars. IRBB55 (a near-isogenic line of IR24), possessing two resistance genes *xa13*/*SWEET11a* and *Xa21*, was selected as a donor parent to introgress these two resistance genes (Gopalakrishnan et al. 2008) in Komboka, FARO-44, and NERICA-4 through marker-assisted backcross breeding (Sundaram et al. 2008; Arra et al. 2024). All the seeds were obtained through a standard material transfer agreement (SMTA) from the International Rice Research Institute, Philippines (IRRI) or AfricaRice (https://www.africarice.org).

### Plant growth conditions

Rice seed germination and plant cultivation were performed as described previously (Luu et al. 2020; Schepler-Luu et al. 2023). Briefly, rice seeds were sterilized and germinated on solid ½ strength Murashige Skoog media (Duchefa, M0222.0050), supplemented with 1% sucrose (Sigma-Aldrich, S7903-250G). Seedlings were grown for 10 days in Magenta boxes before being transferred to soil. Plants were grown in greenhouses at HHU (8-hour day, 30 °C / 16-hour night, 25 °C, relative humidity (RH) 70%, supplemental LED (Valoya, BX100 NS1) at 400 μmol/m^-2^s^-1^). Plants were fertilized weekly from the 2^nd^ week and biweekly from the 6^th^ week after germination (ICL, Peters Excel, CalMag grower, 2152.02.15EB). At IRD, seeds were sown in soil supplemented with 3 g of standard NPK fertilizer (N: 19%; P: 5%; K: 8% per liter) and grown in greenhouses (12 hour day 28 °C, 80% RH, 12 hour night at 25 °C, 60% RH at 200 μmol/m^-2^s^-1^)(Schepler-Luu et al. 2023).

### Generation of Komboka homozygous BC_2_F_3_ plants by MABB

Introgression of two BB-resistance genes, *xa13* and *Xa21,* into the African rice cultivar Komboka was performed by crossing Komboka (*Xa1*+*Xa4*) with IRBB55 (*xa13*+*Xa21*) to generate F_1_ seeds. Heterozygous F_1_ plants for two BB resistance genes*, xa13 and Xa21,* were identified using the foreground gene-linked markers, Xa13prom for *xa13,* and pTA248 for *Xa21*. Selected true single F_1_ plant was backcrossed with the Komboka to generate BC_1_F_1_ seeds, which were subjected to foreground selection using the gene-specific markers for *xa13* and *Xa21*. A single BC_1_F_1_ plant, which is heterozygous for two BB resistance genes, *xa13* and *Xa21*, was then backcrossed with Komboka to generate BC_2_F_1_ seeds. BC_2_F_1_ plants were heterozygous for *xa13* and *Xa21*, as shown by foreground selection and advanced by self-pollination to generate BC_2_F_2_ seeds. Among the BC_2_F_2_ plants, the homozygous plants for four gene combinations (*Xa1*+*Xa4*+*xa13*+*Xa21*) and three gene combinations (*Xa1*+*Xa4*+*xa13*) were identified through foreground selection (Figures 4, 5; Table 1). These BC_2_F_2_ plants were subjected to background selection to identify plants with maximum recipient parent genome recovery. Selected BC_2_F_2_ homozygous four gene combinations (*Xa1*+*Xa4*+*xa13*+*Xa21*) and three gene combinations (*Xa1*+*Xa4*+*xa13*) plants with maximum recipient genome recovery were advanced to the BC_2_F_3_ stage for disease resistance and agromorphological character evaluation.

### Generation of homozygous BC_2_F_3_ inbred lines of FARO-44

A heterozygous F_1_ plant derived from the cross Komboka/ IRBB-55, was used as a donor to attempt crosses with FARO-44 and generate three-way cross F_1_ (TWF_1_) plants. Deployed the foreground selection with gene-specific molecular markers to identify the resistance genes, *Xa4*, *xa13*, and *Xa21,* in the heterozygous stage. One heterozygous F_1_ plant was used as a donor, and attempted backcrosses with FARO-44 were performed to generate BC_1_F_1_ to BC_2_F_1_. Three genes with heterozygous plants were identified and advanced them to the BC_2_F_2_ generation to identify homozygous plants with four resistance genes (*Xa1*, *Xa4*, *xa13*, and *Xa21*). Fore-ground selection was deployed in each generation for the targeted resistance genes (Figure 6a). Homozygous plants with different gene combinations were selected and advanced to the BC_2_F_3_ generation for disease resistance and agronomic and morphological characters.

### Generation of NERICA-4 homozygous BC_1_F_4_ plants by MABB

An F_1_ plant heterozygous for two BB resistance genes (*xa13* and *Xa21*) was identified using foreground gene-linked markers from the Komboka (*Xa1*+*Xa4*)/IRBB55 (*xa13*+*Xa21*) cross was used as a donor to introgress three BB resistance genes (*Xa1*, *xa13*, and *Xa21*) in NERICA-4 by MABB. A cross was attempted between NERICA-4 and the F_1_ plant to generate TWF_1_s. These TWF_1_ plants were screened for three BB resistance genes in a heterozygous state with gene-specific molecular markers, XaL-F1R1 for *Xa1*, Xa13prom for *xa13*, and pTA248 for *Xa21* (Table S4). Selected true TWF_1_ plants were backcrossed to NERICA-4 to generate BC_1_F_1_s, which were subjected to foreground selection using the gene-specific markers. The BC_1_F_1_ plants heterozygous for the *Xa1*, *xa13,* and *Xa21* genes were then self-pollinated to generate BC_1_F_2_ seeds (Figure 6b, Table 1). Among the BC_1_F_2_ plants, the four gene combination (*Xa1*+*Xa4*+*xa13*+*Xa21*) and three gene combination (*Xa1*+*Xa4*+*xa13*) homozygous plants were identified through foreground selection. These BC_1_F_2_ plants were subjected to background selection to identify plants with maximum recipient-parent genome recovery and were advanced to the BC1F4 generation for disease resistance and agronomic-morphological character evaluation.

### Foreground selection

The genomic DNA was isolated from two parents and backcross-derived plants using the peqGOLD Plant DNA Mini Kit (PeqGold, VWR International GmbH, Darmstadt, Germany). The presence or absence of the four target BB resistance genes, i.e., *Xa1*, *Xa4*, *xa13,* and *Xa21,* was confirmed by gene-specific markers, XaL-F1R1 for *Xa1* (Ji et al. 2020), MP1/MP2 for *Xa4* (Sun et al. 2003), xa13-prom for *xa13* (Hajira et al. 2016), and pTA248 for *Xa21* (Table S1)(Ronald et al. 1992). Foreground selection was applied to all generations from F_1_ to BC_x_F_x_ generations to select lines carrying all four target BB resistance genes. We confirmed the hetero or homozygous in BC_2_F_1_s of Komboka and FARO-44, and BC_1_F_1_s of NERICA-4.

### Background selection

SSR molecular markers polymorphic between the donor parent, IRBB55 (*xa13*+*Xa21*), and the recipient parents, Komboka (*Xa1*+*Xa4*), FARO-44, NERICA-4, were screened for a total of 426 rice SSR markers distributed on the rice 12 chromosomes. The polymorphic markers identified between the parents were deployed to identify plants with high RPG recovery in introgressed lines of Komboka (Table S2-S4). To identify the back-cross breeding lines that contain the maximum recovery of RPG (Kumar et al. 2023), a set of polymorphic SSR marker data was used to generate a representative map showing the donor and recipient parents genomic contribution using Graphical Genotype (GGT) software, version 2.0 (Van Berloo 1999).

### Bacterial blight resistance evaluation

Generated backcross-derived breeding lines of Komboka, FARO-44, and NERICA-4 with targeted four gene combinations (*Xa1*+*Xa4*+*xa13*+*Xa21*), three gene combinations *(Xa1+Xa4*+*xa13*), and (*Xa1*+*Xa4*+*Xa21*) were evaluated for BB resistance against a set of representative virulent *Xoo* strains from Tanzania and Madagascar at IRD PHIM Montpellier, France, and HHU, Germany. Selected breeding lines of Komboka, FARO-44, and NERICA-4 were grown in the greenhouse under controlled conditions, and plants at the maximum tillering stage (40-45 days) were leaf clip inoculated with iTz, iMg, and Asian *Xoo* strains. *Xoo* strains were collected from -80 ^°^C stocks and streaked on peptone sucrose agar (PSA) and grown for four days at 28 ^°^C. Single colonies were collected from each strain, plated on the PSA, and incubated for three days at 28 ^°^C. Bacteria were collected from the PSA petri dishes and diluted in sterile water to an OD_600_ of 0.2 for clip-inoculation. Lesion lengths were recorded 14-21 days post-inoculation (Arra et al. 2017; Eom et al. 2019; Oliva et al. 2019; Schepler-Luu et al. 2023).

## Acknowledgments

We thank Gopala Krishnan (ICAR; New Delhi) for critical reading and suggestions on the draft manuscript. This work was supported by Alexander von Humboldt Professorship (WBF), Deutsche Forschungsgemeinschaft (DFG, German Research Foundation), Reinhard Koselleck project (FR 989/14-1); Collaborative Research Center SFB1535, project ID 458090666/CRC1535/1 and Germanýs Excellence Strategy – EXC-2048/1 – project ID 390686111 (CEPLAS), and fellowships to YA by the Alexander von Humboldt Foundation, Department of Biotechnology, India (BT/RLF/Re-entry/48/2022), and support from the Director, CSIR-Central Institute of Medicinal and Aromatic Plants, India.

## Conflict of interest

The authors declare no conflict of interest. Access to lines under MTA from HHU.

## Authors contributions

YA, BS, and WBF developed the concept. YA generated the breeding material by crossing, phenotyping, fore-and background selection. DBN and MS performed background selection and morphological characters. YA, EL, MS, CB, FA, and ET performed disease resistance assay. EL, YA, BS, and WBF wrote and EL and WBF revised the manuscript. All authors have approved the final version of the manuscript.

## Abbreviations

MABB: marker-assisted backcross breeding
iMg: *Xoo* strain introduced into Madagascar
iTz: *Xoo* strain introduced into Tanzania
SSR: marker

## Notes

### Competing Interest Statement

The authors have declared no competing interest.

