## supplementary for "Pyramiding four genes governing bacterial blight resistance in three popular rice varieties of East Africa and Madagascar"

**Table S1.** List of gene-specific primers used for foreground selection

| R-Gene | Primer | Primer sequences ('5-3') | Expected amplicon size | Reference |
|---|---|---|---|---|
| *Xa1* | XaL-F1 | ATCAGGAACTTGAACTCCAG | 270 bp | (Ji et al. 2020) |
| | XaL-R1 | AACCACTGATTGCGGAAGG | | |
| *Xa4* | MP-1F | ATCGATCGATCTTCACGAGG | 180 bp | |
| | MP-2R | TGCTATAAAAGGCATTCGG | | (Sun et al. 2003) |
| *xa13* | xa13 prom-F | GGCCATGGCTCAGTGTTTAT | 488 bp | (Hajira et al. 2016) |
| | xa13 prom-R | GAGCTCCAGCTCTCCAAATG | | |
| *Xa21* | pTA248_F | AGACGCGGAAGGGTGGTTCCCGGA | 960 bp | (Ronald et al. 1992) |
| | pTA248_R | AGACGCGGTAATCGAAAGATGAAA | | |

**Table S2.** List of SSR markers used for background selection of Komboka x IRBB55 cross

| Chromosome | Screened SSRs | Polymorphic SSRs | List of polymorphic SSRs | Target *Xa* Gene |
|---|---|---|---|---|
| 1 | 31 | 6 | RM246, RM572, RM579, RM580, RM594, RM3412 | - |
| 2 | 24 | 2 | RM110, RM154 | - |
| 3 | 18 | 2 | RM489, RM5474 | - |
| **4** | **71** | **12** | **RM6314, RM16649, RM16653, RM16569, RM16790, RM16801, RM16815, RM16817, RM16819, RM16821, RM16913, RM17093** | *Xa1* |
| 5 | 25 | 8 | RM163, RM274, RM413, RM1237, RM2998, RM5579, RM7446, RM18614 | - |
| 6 | 28 | 2 | RM444, RM19290 | - |
| 7 | 21 | 3 | RM180, RM8007, RM21260 | - |
| **8** | **70** | **17** | **RM152, RM447, RM1345, RM3496, RM3754 RM3819, RM5493, RM6925, RM10383, RM22529, RM22539, RM22540, RM22837, RM23523, RM23386, RM23502, RM 25503** | *xa13* |
| 9 | 27 | 5 | RM434, RM2393, RM23872, RM23937, RM23948 | - |
| 10 | 18 | 7 | RM171, RM216, RM271, RM1375, RM1873, RM5147, RM25262 | - |
| **11** | **78** | **18** | **RM202, RM206, RM555, RM2191, RM3717, RM25982, RM26002, RM26063, RM26352, RM26402, RM26524, RM26799, RM26823, RM26956, RM26957, RM26931, RM26969, RM27367/144** | *Xa4* **and** *Xa21* |
| 12 | 15 | 3 | RM19, RM235, RM3331 | - |
| Total SSRs | **426** | **85** | | |

**Note:** Markers highlighted in bold represent the target genes, *Xa1*, *Xa4*, *xa13*, and *Xa21*. **-**: Non-targeted *Xa* gene.

**Table S3.** List of SSR markers used for background selection of FARO-44 x Komboka/IRBB55 cross

| **Chromosome** | **Screened SSRs** | **No of polymorphic SSRs** | **Polymorphic SSRs** | **Target *Xa* gene** |
|---|---|---|---|---|
| 1 | 31 | 0 | 0 | - |
| 2 | 24 | 2 | RM71, RM1385 | - |
| 3 | 18 | 1 | RM520 | - |
| **4** | **71** | **7** | **RM5687, RM16649, RM16653, RM16657, RM16819, RM16821, RM16824** | ***Xa1*** |
| 5 | 25 | 1 | RM248 | - |
| 6 | 28 | 1 | RM19290 | - |
| 7 | 21 | 1 | RM21591 | - |
| **8** | **70** | **13** | **RM547, RM10383, RM22692, RM22529, RM22533, RM22541, RM22544, RM22548, RM22550, RM22551, RM22552, RM22555, RM22556** | ***xa13*** |
| 9 | 27 | 1 | RM24017 | - |
| 10 | 18 | 1 | RM258 | - |
| **11** | **78** | **15** | **RM1219, RM3754, RM4469, RM6085, RM6690, RM25951, RM26158, RM26334, RM26513, RM26666, RM26669, RM26937, RM26945, RM26956, RM26957** | ***Xa4,* and *Xa21*** |
| 12 | 15 | 0 | 0 | - |
| Total SSRs | 426 | 43 | | |

**Note:** Markers highlighted in bold represent the target genes, *Xa1*, *Xa4*, *xa13*, and *Xa21*.-: Non-targeted *Xa* gene.

**Table S4.** List of SSR markers used for background selection of NERICA-4 x Komboka/IRBB55 cross

| **Chromosome** | **Screened SSRs** | **Polymorphic SSRs** | **List of polymorphic SSRs** | **Target *Xa* Gene** |
|---|---|---|---|---|
| 1 | 31 | 3 | RM151, RM493, RM580 | - |
| 2 | 24 | 4 | RM424, RM526, RM1385, RM7581 | - |
| 3 | 18 | 0 | 0 | - |
| **4** | **71** | **5** | **RM16447, RM16569, RM16643, RM17093, RM17515** | ***Xa1*** |
| 5 | 25 | 4 | RM289, RM2998, RM5579, RM5592 | - |
| 6 | 28 | 3 | RM19289, RM19290, RM20068 | - |
| 7 | 21 | 5 | RM248, RM533, RM2752, RM6728, RM8007 | - |
| **8** | **70** | **13** | **RM310, RM547, RM3409, RM3598, 3754, RM5493, RM6925, RM22544, RM22551, RM22552, RM22559, RM23522, RM22837** | ***xa13*** |
| 9 | 27 | 6 | RM201, RM242, RM257, RM2393, RM5777, RM23946 | - |
| 10 | 18 | 6 | RM258, RM311, RM474A, RM1873, RM5147, RM25110 | - |
| **11** | **78** | **19** | **RM229, RM1812, RM4469, RM6085, RM6690, RM26158, RM26187, RM26662, RM26666, RM26334, RM26352, RM26402, RM26438, RM26577, RM26756, RM26830, RM26937, RM26938, RM27367/144** | ***Xa4* and *Xa21*** |
| 12 | 15 | 6 | RM235, RM247, RM1080, RM1226, RM2972, JGT12186 | - |
| Total SSRs | **426** | **74** | | |

**Note:** Markers highlighted in bold represent the target genes, *Xa1*, *Xa4*, *xa13*, and *Xa21*. **-**: Non-targeted *Xa* gene.

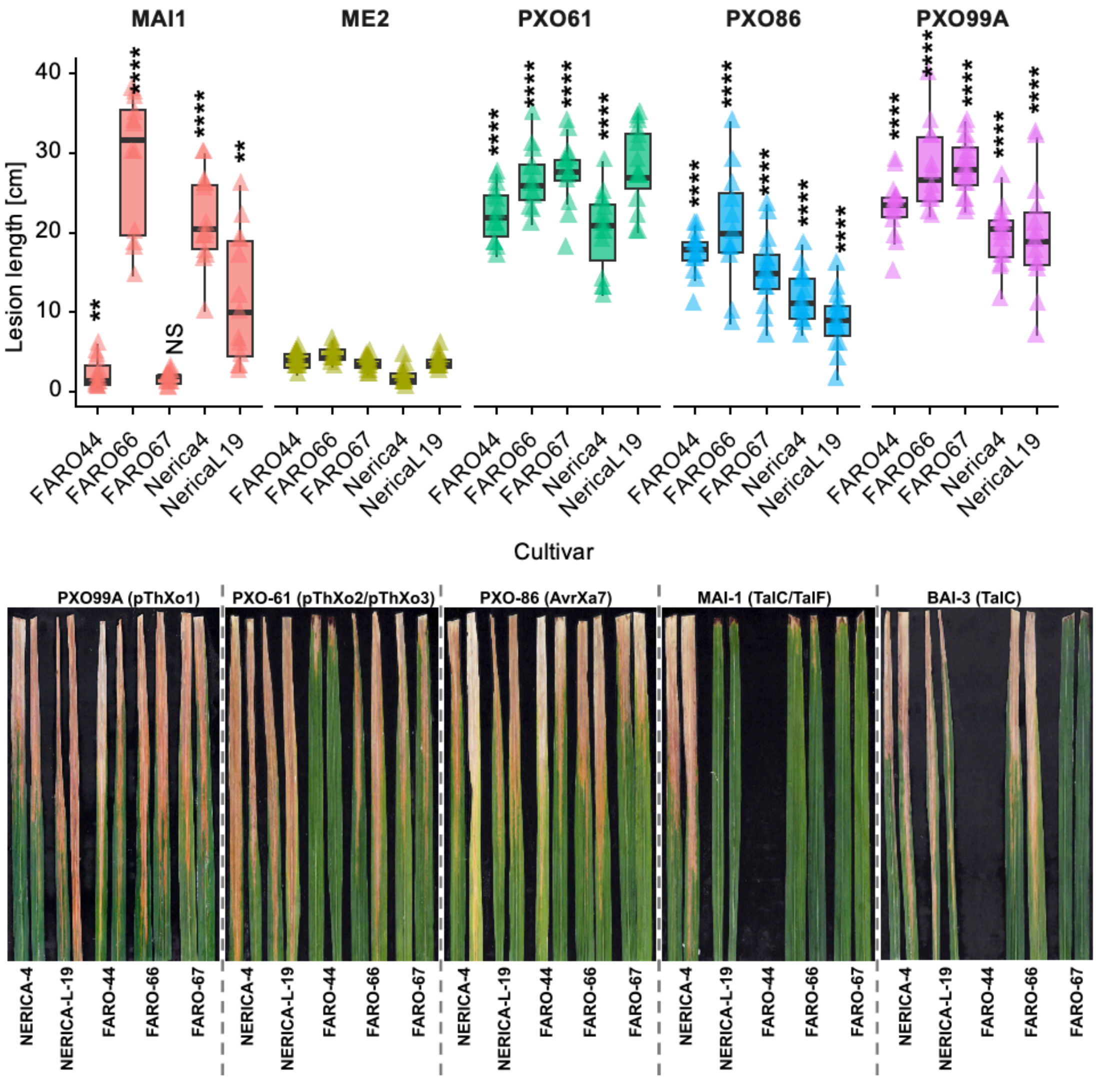


**Figure S1: Virulence of *Xoo* strains carrying major TALes on indicated elite rice variety.**
Top: Independent repeats for Figure 1c. P-values calculated using Student's T-test against ME2 of the respective cultivar. NS- not significant, ** - 0.01, **** - 0.0001. N > 10, assay repeated three time with comparable results. Bottom: Phenotypic characterization of lesion length observed in indicated elite rice cultivars 14-days post-infection with indicated *Xoo* strains.

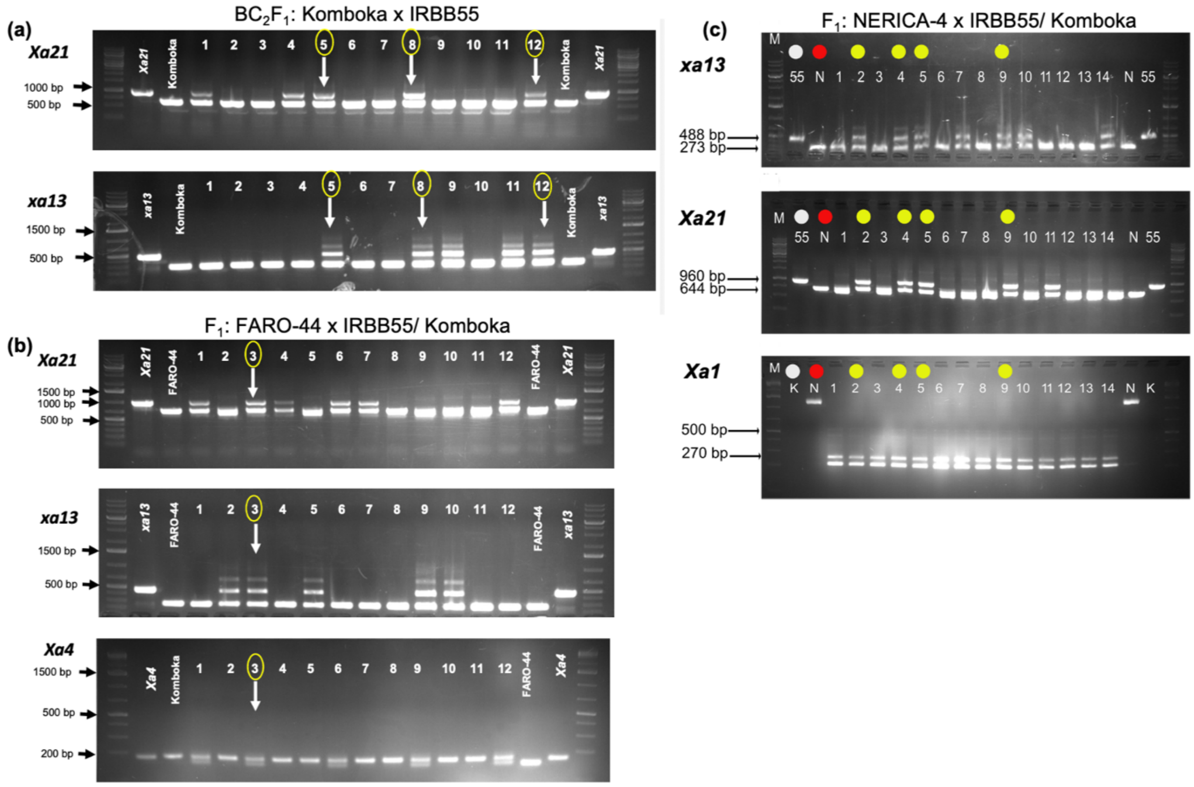


**Figure S2: Foreground selection for *Xa* genes in plants derived from MABB.**
Genotyping for indicated *Xa* genes in progenies derived from a) IRBB55 x Komboka; b) FARO-44 x IRBB55/Komboka; and c) NERICA-4 x IRBB55/Komboka. White circles indicate donor parent, red circles indicate recipient parent, and indicated with a yellow circle are selected progenies used for advanced crosses. K- Komboka, N- NERICA-4 (related to Figure 4).

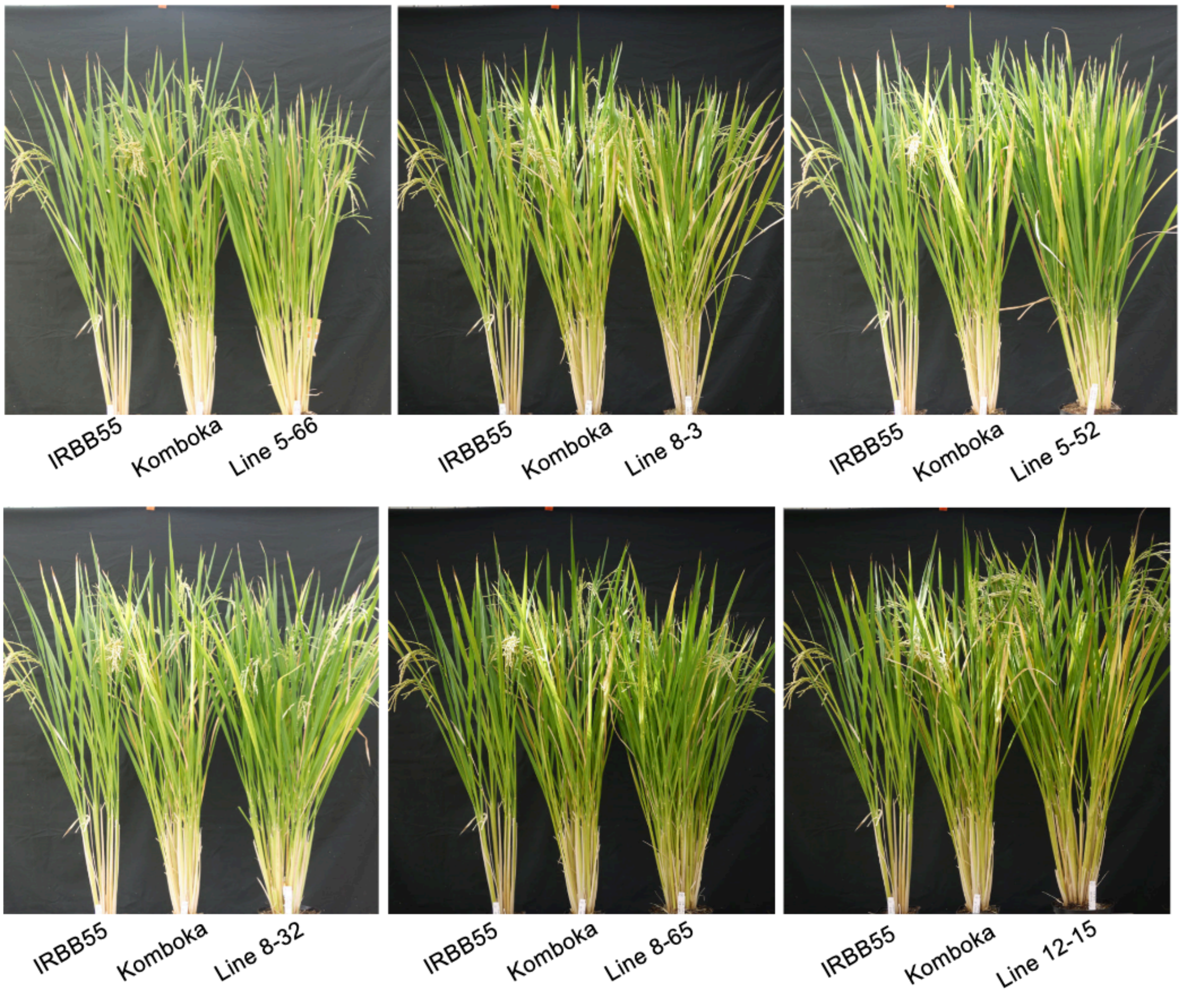


**Figure S3: Phenotype of six independent Komboka x IRBB55 ($BC_2F_2$) crossing lines after seed setting (homozygous for all four R-genes: *Xa1*, *Xa4*, *xa13*, and *Xa21*).** Crossing lines are indicated at the bottom of each image.

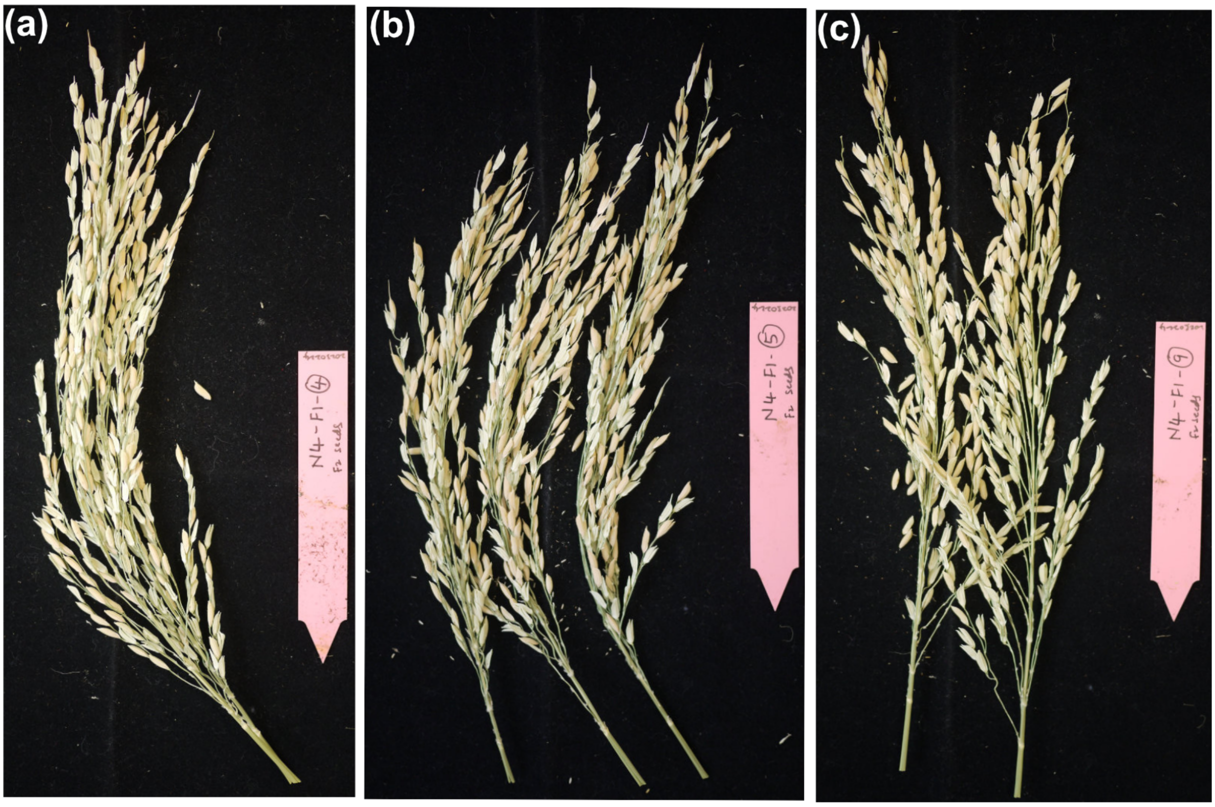


**Figure S4: Seed setting in panicles $F_1$ plants derived from NERICA-4 x IRBB55/Komboka.**
(a) $F_2$ seeds from MABB line 4; (b) $F_2$ seeds from MABB line 5; (c) $F_2$ seeds from MABB line 9.

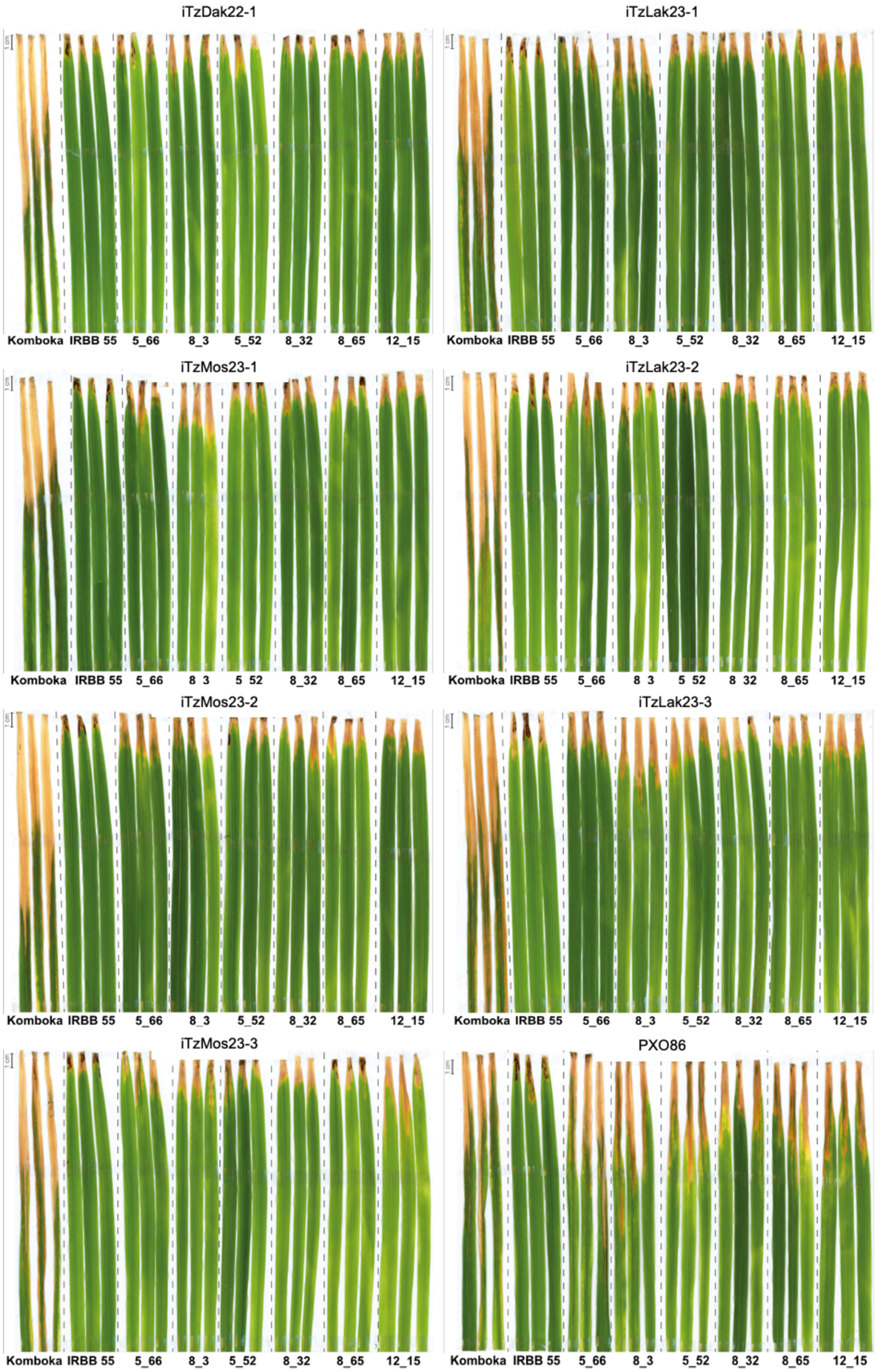


**Figure S5: Disease symptoms of $BC_2F_3$ Komboka/IRBB55 MABB lines clip-infected with iTz and iMg *Xoo* strains (cf. also Table S2).** Individual plants at 40-45 days were clip-inoculated with representative *Xoo* strains. Phenotypic observations were recorded by measuring the lesion length (cm) 21 days post-infection (related to Figure 7).

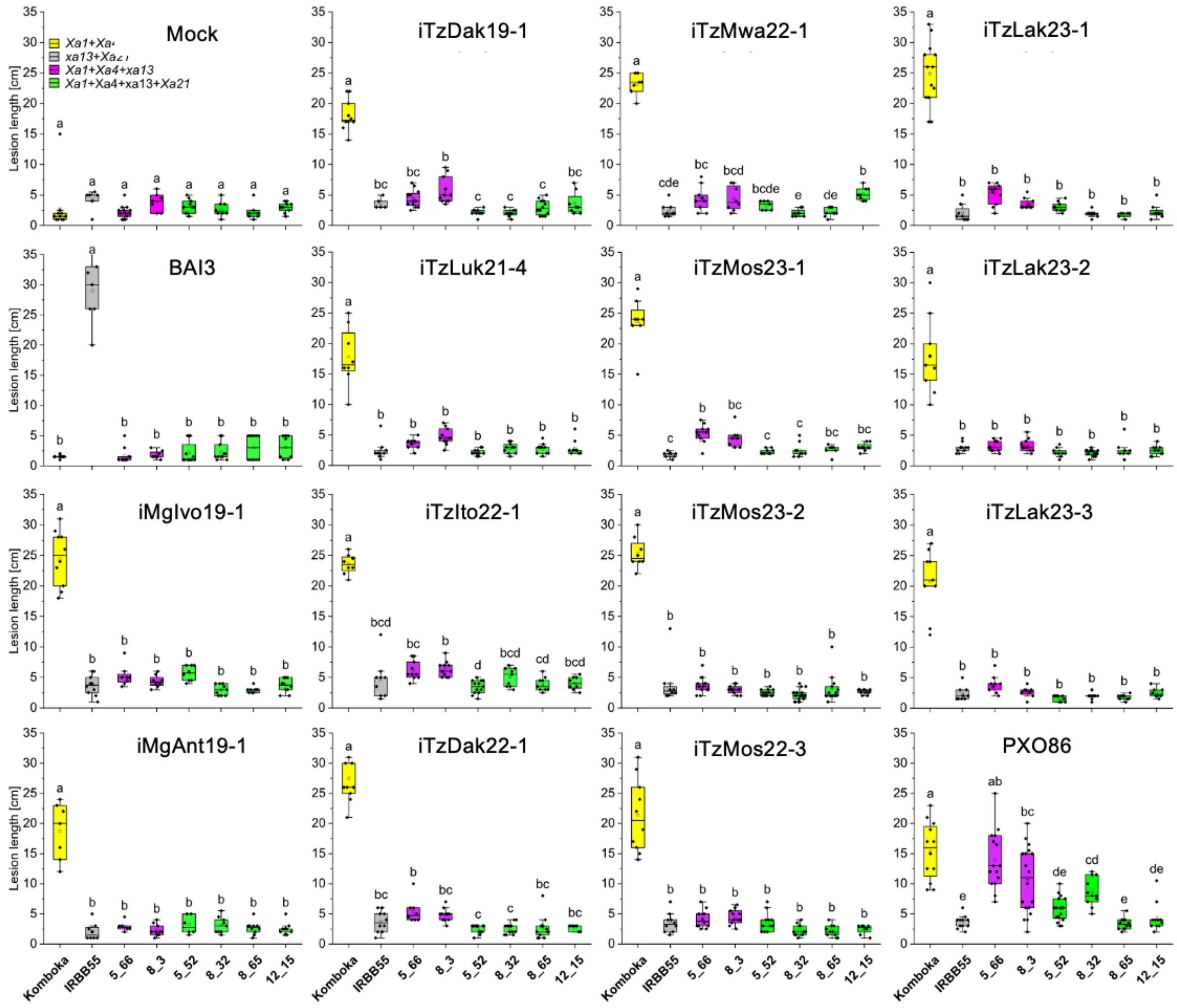


**Figure S6: Virulence profile of *Xoo* strains from Tanzania and Madagascar on MABB Komboka breeding lines.** Individual $BC_2F_3$ generation breeding lines and parents were inoculated with indicated iTz and iMg *Xoo* strains. BB symptoms were recorded 21 days post-inoculation by measuring the lesion length (cm). Boxes extend from 25th to 75$^{th}$ percentiles and display median values as center lines. Whiskers mark the minimum and maximum values; asterisks indicate each data points. Significance was evaluated by one-way ANOVA followed by the Bonferroni test; different letters indicate significant differences ($P < 0.05$) (related to Figure 7).

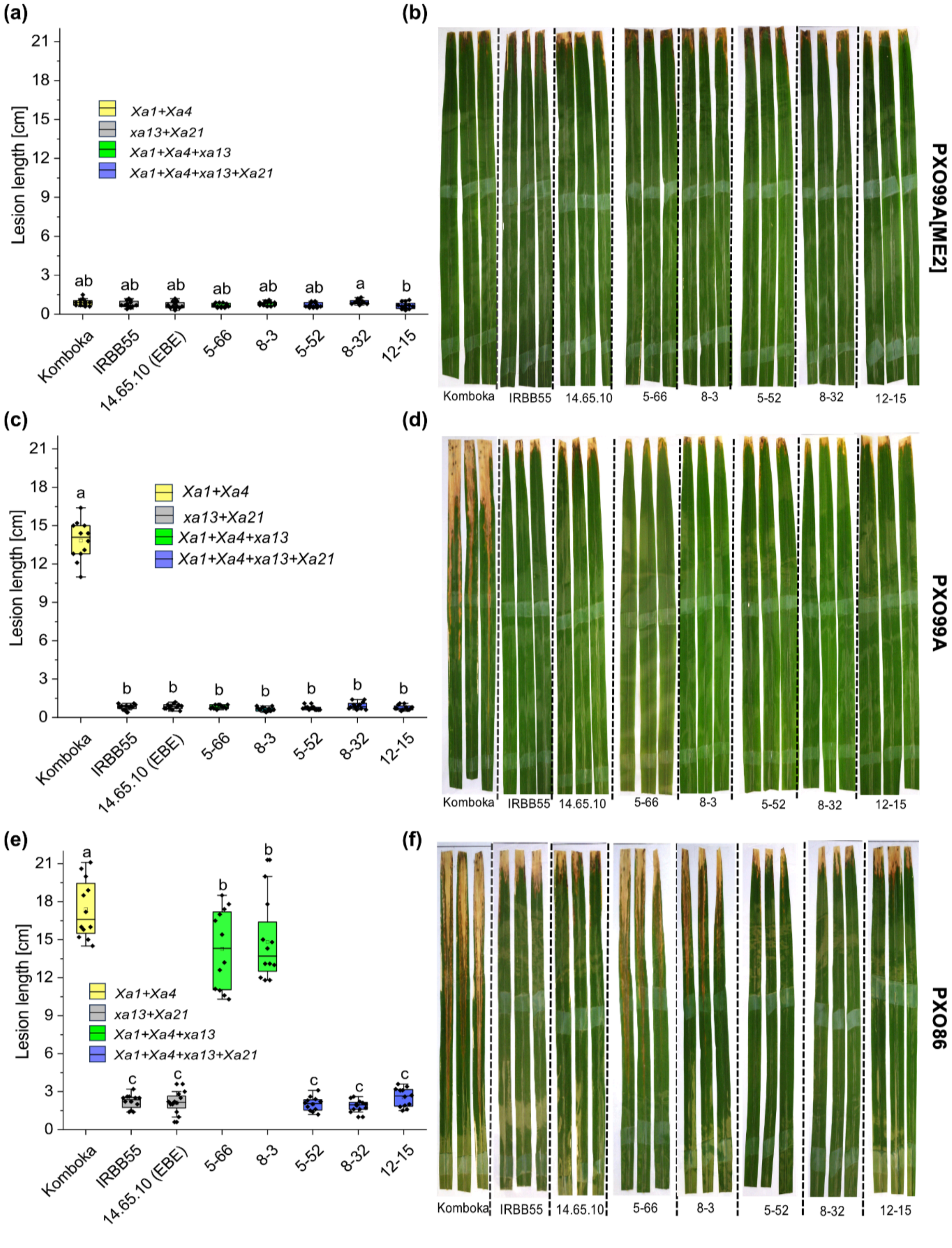


**Figure S7: Disease symptoms of individual $BC_2F_3$ Komboka/IRBB55 MABB lines clip-infected with Asian *Xoo* strains**. Lesion length quantification and their corresponding phenotype of individual 40-45 day-old plants 21 days post-infectinon with indicated *Xoo* strains. Lesion length and phenotype for indicated lines leaf clip-inoculated with **(a-b)** avirulent PXO99A[ME2], **(c-d)** PXO99A carrying pThXo1 targeting *SWEET11a,* and **(e-f)** PXO86 carrying AvrXa7 targeting *SWEET14.* 14.65.10 (EBE): Genome-edited Komboka line with modifications in the pThXo1, pThXo2, pThXo3, TalC, AvrXa7, and TalF EBEs (Schepler-Luu et al. 2023) (related to Figure 7).
